# Epitope-Based Peptide Vaccine Against Severe Acute Respiratory Syndrome-Coronavirus-2 Nucleocapsid Protein: An *in silico* Approach

**DOI:** 10.1101/2020.05.16.100206

**Authors:** Ahmed Rakib, Saad Ahmed Sami, Md. Ashiqul Islam, Shahriar Ahmed, Farhana Binta Faiz, Bibi Humayra Khanam, Mir Muhammad Nasir Uddin, Talha Bin Emran

**Affiliations:** Department of Pharmacy, Faculty of Biological Sciences, University of Chittagong, Chittagong-4331, Bangladesh; Department of Pharmacy, Mawlana Bhashani Science & Technology University, Santosh, Tangail-1902, Bangladesh; Drug Discovery, GUSTO A Research Group, Chittagong-4000, Bangladesh; Department of Pharmacy, BGC Trust University Bangladesh, Chittagong-4381, Bangladesh

**Keywords:** COVID-19, SARS-CoV-2, de novo vaccine, epitope, immunity

## Abstract

With an increasing fatality rate, severe acute respiratory syndrome-coronavirus-2 (SARS-CoV-2) has emerged as a promising threat to human health worldwide. SARS-CoV-2 is a member of the *Coronaviridae* family, which is transmitted from animal to human and because of being contagious, further it transmitted human to human. Recently, the World Health Organization (WHO) has announced the infectious disease caused by SARS-CoV-2, which is known as coronavirus disease-2019 (COVID-2019) as a global pandemic. But, no specific medications are available for the treatment of COVID-19 so far. As a corollary, there is a need for a potential vaccine to impede the progression of the disease. Lately, it has been documented that the nucleocapsid (N) protein of SARS-CoV-2 is responsible for viral replication as well as interferes with host immune responses. We have comparatively analyzed the sequences of N protein of SARS-CoV-2 for the identification of core attributes and analyzed the ancestry through phylogenetic analysis. Subsequently, we have predicted the most immunogenic epitope for T-cell as well as B-cell. Importantly, our investigation mainly focused on major histocompatibility complex (MHC) class I potential peptides and NTASWFTAL interacted with most human leukocyte antigen (HLA) that are encoded by MHC class I molecules. Further, molecular docking analysis unveiled that NTASWFTAL possessed a greater affinity towards HLA and also available in a greater range of the population. Our study provides a consolidated base for vaccine design and we hope that this computational analysis will pave the way for designing novel vaccine candidates.

## 1 Introduction

The present world has witnessed the outbreak of many life-threatening human pathogens including Ebola, Chikungunya, Zika, Severe Acute respiratory syndrome coronavirus (SARS-CoV), Middle East respiratory syndrome coronavirus (MERS-CoV) in the 21^st^ century. More recently in late December 2019, a cluster of pneumonia cases was reported in the city of Wuhan, Hubei province, China which was of unknown cause. Later it was confirmed that these pneumonia cases were due to a novel coronavirus named SARS-CoV-2 (previously named as 2019-nCoV) and the disease condition of this virus is referred to as COVID-19 (1–3). On March 11, 2020, the World Health Organization (WHO) has assessed that COVID-19 can be characterized as a pandemic. The current COVID-19 pandemic is a global concern and is spreading at an alarming rate and as of April 12, 2020, more than 1.6 million cases and over 105,000 deaths have been reported globally (4).

Coronaviruses (CoVs) are phenotypically and genotypically diverse group viruses that can adapt to the new environment through mutation and recombination, probably even more than influenza. Coronaviruses often infect mammals, birds and can transmit to humans. Six strains of coronaviruses were found in the last few decades but this is a completely new strain and of zoonotic origin. COVID-19 virus belongs to the *Coronaviridae* family of the Genus *Betacoronavirus*, pleomorphic or spherical particles, 150 to 160 nm in size, associated with positive single-strand RNA (ssRNA) which is surrounded by crown-shaped, club-like spikes projection on the outer surface. Among all RNA viruses, Coronaviruses have the largest genome typically ranging from 27 to 32 kb. After the two previously reported coronavirus-SARS-CoV and MERS-CoV, this is the third coronavirus that has already infected humans and the preliminary investigations revealed that some environmental specimens of the Huanan seafood market in Wuhan were positive for COVID-19 (3). Although the seafood market was reckoned positive for COVID-19, no specific association with an animal is confirmed yet based on the WHO report. Researchers are working to establish a possible animal reservoir for COVID-19 (5).

As COVID-19 is mainly a respiratory disease, in most cases it might affect the lungs only. The primary mode of infection is human-to-human transmission through close contact, which occurs via spraying droplets from the infected individual through their cough or sneeze. The symptoms of this coronavirus can be mild to moderate or severe including, fever, cough, and shortness of breath or pneumonia. Respiratory, hepatic and neurological complications can be seen in case of severe cases that can lead to death. It seems that the severity and fatality rate of COVID-19 is milder than that of SARS and MERS. Although diarrhea was presented in about 20-25% of patients with SARS and MERS, intestinal symptoms were rarely reported in patients with COVID-19 (6–8). Multi-organ failure, especially in elderly people and people with underlying health conditions such as hypertension, cardiovascular disease and diabetes, are exhibiting a higher mortality rate in COVID-19.

Interestingly, SARS-CoV-2 has 82% similarity with the original SARS-CoV virus attributed to the outbreak in 2003 (9). A mature SARS-CoV-2 virus generally has a polyprotein (the open reading frame 1a and 1b, Orf1ab), four structural proteins such as envelope (E) protein; membrane (M) protein; nucleocapsid (N) protein; spike (S) protein and five accessory proteins (Orf3a, Orf6, Orf7a, Orf8, Orf10), and particularly, SARS-CoV-2 encodes an additional glycoprotein having acetyl esterase and hemagglutination (HE) attributes, which identified it distinct than its two predecessors (10). The functions of accessory proteins may include signal inhibition, apoptosis induction and cell cycle arrest (11). The S protein on the surface of the viral particle enables the infection of host cells by binding to the host cell receptor angiotensin-converting enzyme 2 (ACE2), utilizing the S-protein’s receptor-binding domain (RBD).

The N protein binds to the RNA genome of the COVID-19 and creates a shell or capsid around the enclosed nucleic acid. N protein is involved in viral RNA synthesis and folding which interacts with the viral membrane protein during viral assembly affects host cell responses including cell cycle and translation. An epitope-based peptide vaccine has been raised in this aspect. The core mechanism of the peptide vaccine is based on the chemical method to synthesize the recognized B-cell and T-cell epitopes that can induce specific immune responses and are immune-dominant. T-cell epitopes are short peptide fragments (8-20 amino acids) while the B-cell epitopes can be proteins (12,13).

Once a mutated virus infects the host cells by escaping the antibodies, it then relies upon the T-cell mediated immunity to fight against the virus. Viral proteins are processed into short peptides inside the infected cells and then loaded onto major histocompatibility complexes (MHC) proteins. After that, the MHC-peptide complexes are presented on the infected cell surface for recognition by specific T cells. Activated CD8^+^ T cells then recognize the infected cells and clear them. T-cell immunity also depends strictly on the MHC-peptide complexes which are similar to the antigen-antibody association. MHC proteins are encoded by human leukocyte antigen (HLA) which is located among the most genetically variable regions on the human genome. Each HLA allele can only present a certain set of peptides that can be presented on the infected cell surface and recognized by T cells are called T-cell epitopes. For a vaccine, it is essential to identify T-cell epitopes that originate from conserved regions of the virus T cell responses against the S and N proteins have been reported to be the most dominant and long-lasting (14).

To develop effective diagnostic tests and vaccine, the identification of B-cell and T-cell epitopes for SARS-CoV-2 proteins are critical especially for structural N and S proteins. Both humoral immunity and cellular immunity provided by B-cell antibodies and T-cells respectively are essential for effective vaccines (15,16). Although humans may mount an antibody response against viruses normally, only neutralizing antibodies can block the entry of viruses into human cells completely (17). Antibody binding site’s location on a viral protein strongly affects the body’s ability to produce neutralizing antibodies(18). It is important to understand whether SARS-CoV-2 has potential antibody binding sites (B-cell epitopes) near their interacting surface with its known human entry receptor, ACE2. Besides neutralizing antibodies, human bodies also depend on cytotoxic CD8^+^ T-cells and helper CD4^+^ T-cells to clear viruses completely from the body. For anti-viral T-cell responses, presentation of viral peptides by human MHC class I and class II is essential (19). MHC-I analysis includes common alleles for HLA-A, HLA-B, and HLA-C. Multiple investigations have indicated that antibodies generated against the N protein of SARS-CoV are highly immunogenic and abundantly expressed protein during infection (20).

The purpose of our present study is to promote the designing of a vaccine against COVID-19 using *in silico* methods, considering SARS-CoV-2 N protein. The reason for focusing particularly on the epitopes in the N structural proteins is due to their dominant and long-lasting immune response which was reported against SARS-CoV previously. For the identified T-cell epitopes, we incorporated the information on the associated MHC alleles so that we can provide a list of epitopes that seek to maximize population coverage globally. Therefore, we designed an epitope-based peptide vaccine to potentially narrow down the search for potent targets against SARS-CoV-2 using the computational approach with an expectation that the wet laboratory research will validate our result.

## 2 Materials and Methods

The methodologies used for peptide vaccine development for SARS-CoV-2 N protein are shown in **Figure 1**.

**FIGURE 1.**
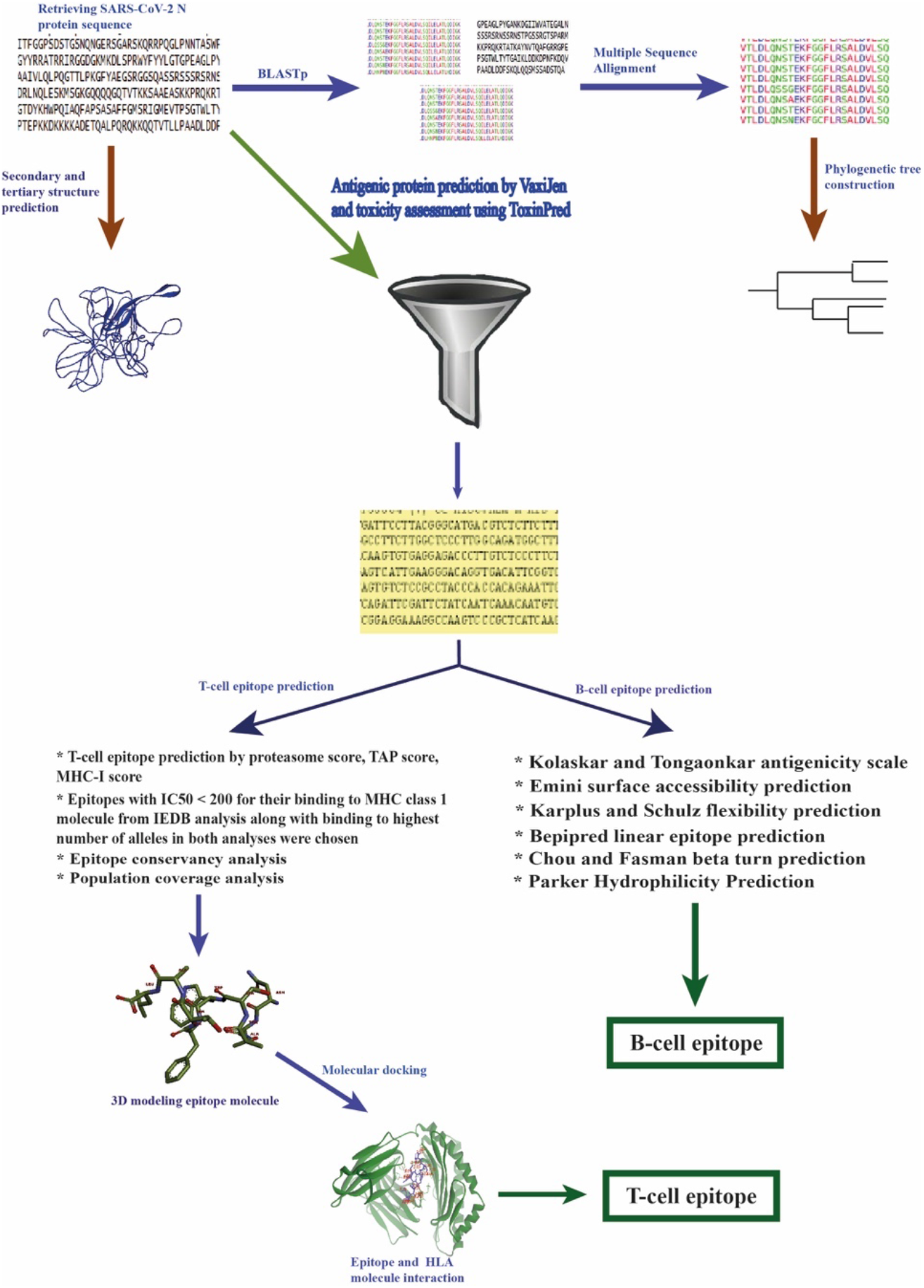
Workflow of the methodologies used in peptide vaccine design.

### 2.1 Protein sequence retrieval

The SARS-CoV-2 N protein sequence was extracted from NCBI (National Center for Biotechnology Information) protein database (Accession no.: QIC53221.1, GI: 1811294683) in FASTA format.

### 2.2 Sequence analysis

The understanding of the features, function, structure, and evaluation is mainly based on the process of sequence analysis which depicts the process of subjecting DNA, RNA, or peptide sequences to wide ranges of analytical methods. We implied NCBI BLAST (Basic Local Alignment Search Tool) (21) that screens homologous sequences from its database and selects those sequences that are more similar to our SARS-CoV-2 N protein; we also performed multiple sequence alignment (MSA) using the ClustalW web server with default settings, and a phylogenetic tree was assembled using MEGA6 software and a web logo was also generated for the conserved peptide sequences using WebLogo based on this alignment (21–23).

### 2.3 Protein antigenicity and toxicity prediction

To determine the potent antigenic protein of the SARS-CoV-2 N protein, we used the online server VaxiJen v2.0, with a default threshold value (24). All the antigenic proteins of SARS-CoV-2 N protein with their respective scores were obtained then sorted in Notepad++. A single antigenic protein with maximum antigenicity scores was selected for further evaluation. The toxicity of epitopes was analyzed using the Toxinpred web server (25).

### 2.4 Protein secondary and tertiary structure prediction

The secondary structure of the SARS-CoV-2 N protein was predicted by using CFSSP (Chou & Fasman Secondary Structure Prediction) because the antigenic part of the protein is more likely to belong to the β-sheet region (26). Also, we predicted the 3D structure of the protein using EasyModeller, a graphical user interface (GUI) version of MODELLER, where we designed the three-dimensional structure of the SARS-CoV-2 N protein using template proteins from Protein Data Bank. The model was validated using PROCHECK and PROSA web servers (27–30).

### 2.5 T-cell epitope prediction

#### 2.5.1 CD8^+^ T-cell epitope prediction

For the *de novo* prediction of T-cell epitope, NetCTL 1.2 server was used in this experiment, using a 0.95 threshold to maintain sensitivity and specificity of 0.90 and 0.95, respectively. The tool expands the prediction for 12 MHC-I supertypes and integrates the prediction of peptide MHC-I binding, proteasomal C-terminal cleavage with TAP transport efficiency. These predictions were performed by an artificial neural network, weighted TAP transport efficiency matrix and a combined algorithm for MHC-I binding and proteasomal cleavage efficiency was then used to determine the overall scores and translated into sensitivity/specificity. Based on this overall score, five best peptides (epitopes) were selected for further evaluation.

For the prediction of peptides binding to MHC-I, we used a tool from the Immune Epitope Database (IEDB) and calculate IC50 values for peptides binding to specific MHC-I molecules (31). For the binding analysis, all the frequently used alleles were selected with a word length of nine residues and binding affinity < 200 nm for further analysis. Another tool (named as MHC-NP) provided by the IEDB server was used to assess the probability that a given peptide was naturally processed and bound to a given MHC molecule (32).

### 2.6 Epitope conservancy and immunogenicity prediction

The degree of similarity between the epitope and the target (i.e. given) sequence is elucidated by epitope conservancy. This property of epitope gives us the promise of its availability in a range of different strains. Hence for the analysis of the epitope conservancy, the web-based tool from IEDB analysis resources was used (33). Immunogenicity prediction can uncover the degree of influence (or efficiency) of the respective epitope to produce an immunogenic response. The T-cell class I pMHC immunogenicity predictor at IEDB, which uses amino acid properties as well as their position within the peptide to predict the immunogenicity of a class I peptide MHC (pMHC) complex (34).

### 2.7 Prediction of population coverage and allergenicity assessment

The population coverage tool from IEDB was applied to determine the population coverage for every single epitope by selecting HLA alleles of the corresponding epitope.

Allergenicity of the predicted epitope was calculated using AllerTop v2.0(35), which is an alignment-free server, used for *in silico* based allergenicity prediction of a protein-based on its physiochemical properties.

### 2.8 HLA and epitope interaction analysis using molecular docking studies

#### 2.8.1 Epitope model generation

The 3D structures of the selected epitopes were predicted by PEP-FOLD, a web-based server (36). For each sequence, the server predicted five probable structures. The energy of each structure was determined by SWISS-PDB VIEWER and the structure with the lowest energy was chosen for further analysis.

#### 2.8.2 Retrieval of HLA allele molecule

The three-dimensional structure of the HLA-A*68:02 (PDB ID: 4I48) was retrieved from Protein Data Bank (RCSB-PDB).

#### 2.8.3 Molecular docking analysis

Molecular docking analysis was performed using Autodock vina in PyRx 0.8, by considering the HLA-A*68:02 molecule as receptor protein and identified epitopes as ligand molecule (37). Firstly, we used the protein preparation wizard of UCSF Chimera (Version 1.11.2) to prepare the protein for docking analysis by deleting the attached ligand, adding hydrogens and Gasteiger–Marsili charges (38,39). The prepared file was then added to the Autodock wizard of PyRx 0.8 and converted into pdbqt format. The energy form of the ligand was minimized and converted to pdbqt format by OpenBabel (40). The parameters used for the docking simulation were set to default. The size of the grid box in AutoDock Vina was kept at 50.183 × 50.183 × 50.183 Å respectively, for X, Y, and Z-axis. AutoDock Vina was implemented via the shell script offered by AutoDock Vina developers. Docking results were observed by the negative score in kcal/mol, as the binding affinity of ligands (41).

### 2.9 B-cell epitope identification

The prediction of B-cell epitopes was performed to find the potential antigen that assures humoral immunity. To detect B-cell epitope, various tools from IEDB were used to identify the B-cell antigenicity, together with the Emini surface accessibility prediction, Kolaskar and Tongaonkar antigenicity scale, Karplus and Schulz flexibility prediction, Bepipred linear epitope prediction analysis and since antigenic parts of a protein belonging to the beta-turn regions, the Chou and Fasman beta-turn prediction tool was also used (42–47).

## 3 Results

### 3.1 Sequence retrieval and analysis

We retrieved the SARS-CoV-2 N protein sequence from the NCBI database (Accession No.: QIC53221.1). Then we performed BLASTp using NCBI-BLAST for the nucleocapsid protein of SARS-CoV-2. We searched for a total of 100 homologs with > 60% identical sequences. Multiple sequence alignment was then performed **(Supplementary data 1)**, and a phylogenetic tree was constructed **(Supplementary Figure S1)**. From multiple sequence alignment, it has been confirmed that the protein sequences have a close relationship. A web logo was designed using the WebLogo server to demonstrate the conserved region **(Figure 2)**.

**FIGURE 2.**
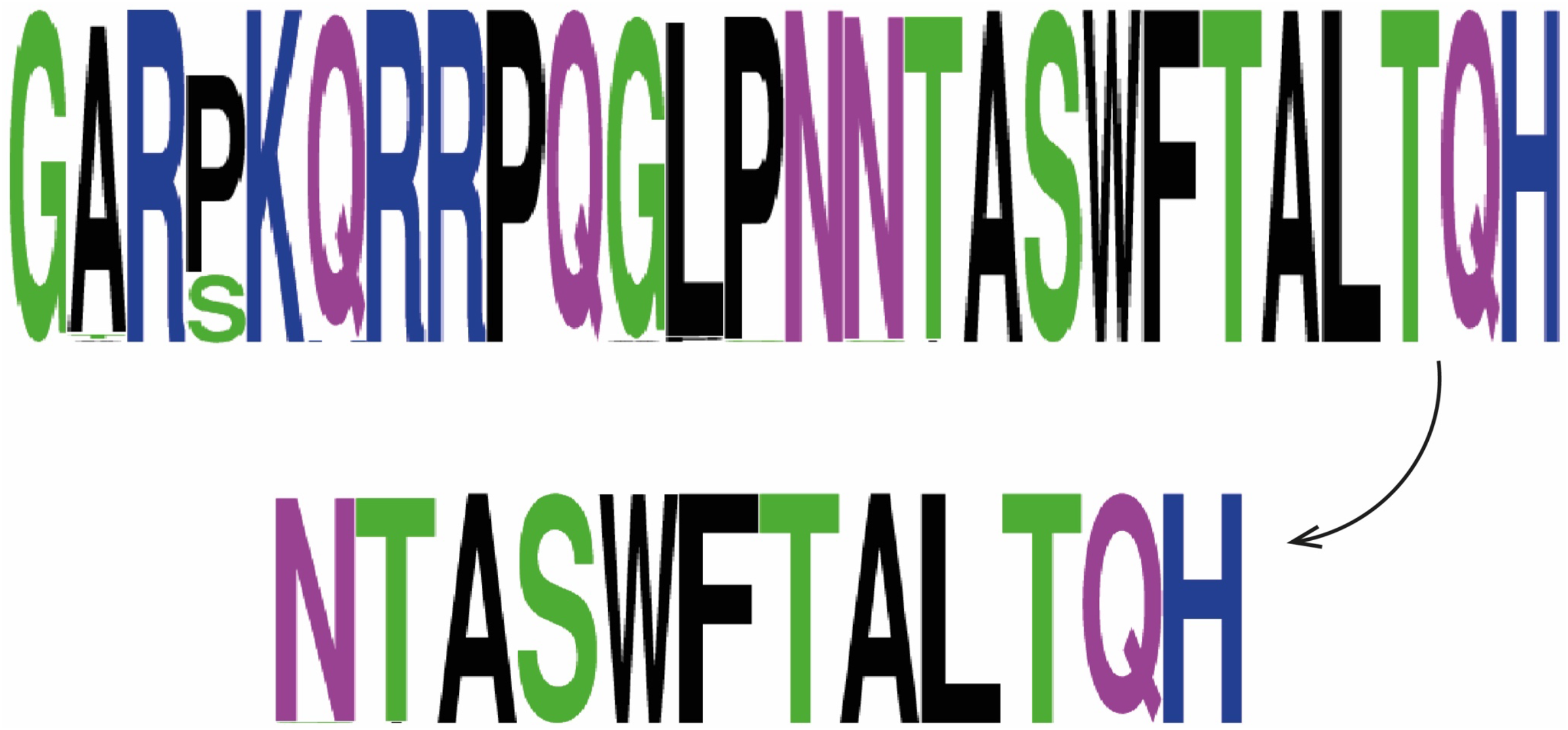
Logo of the conserved peptide sequence.

### 3.2 Antigenic protein prediction

The most potent antigenic protein of SARS-CoV-2 N protein was predicted by VaxiJen v2.0, which is based on the auto-cross covariance transformation of protein sequences into uniform vectors of principal amino acid properties **(Table 1).** The overall antigen prediction score was 0.5002 (probable antigen) at 0.4 threshold value.

**TABLE 1.**
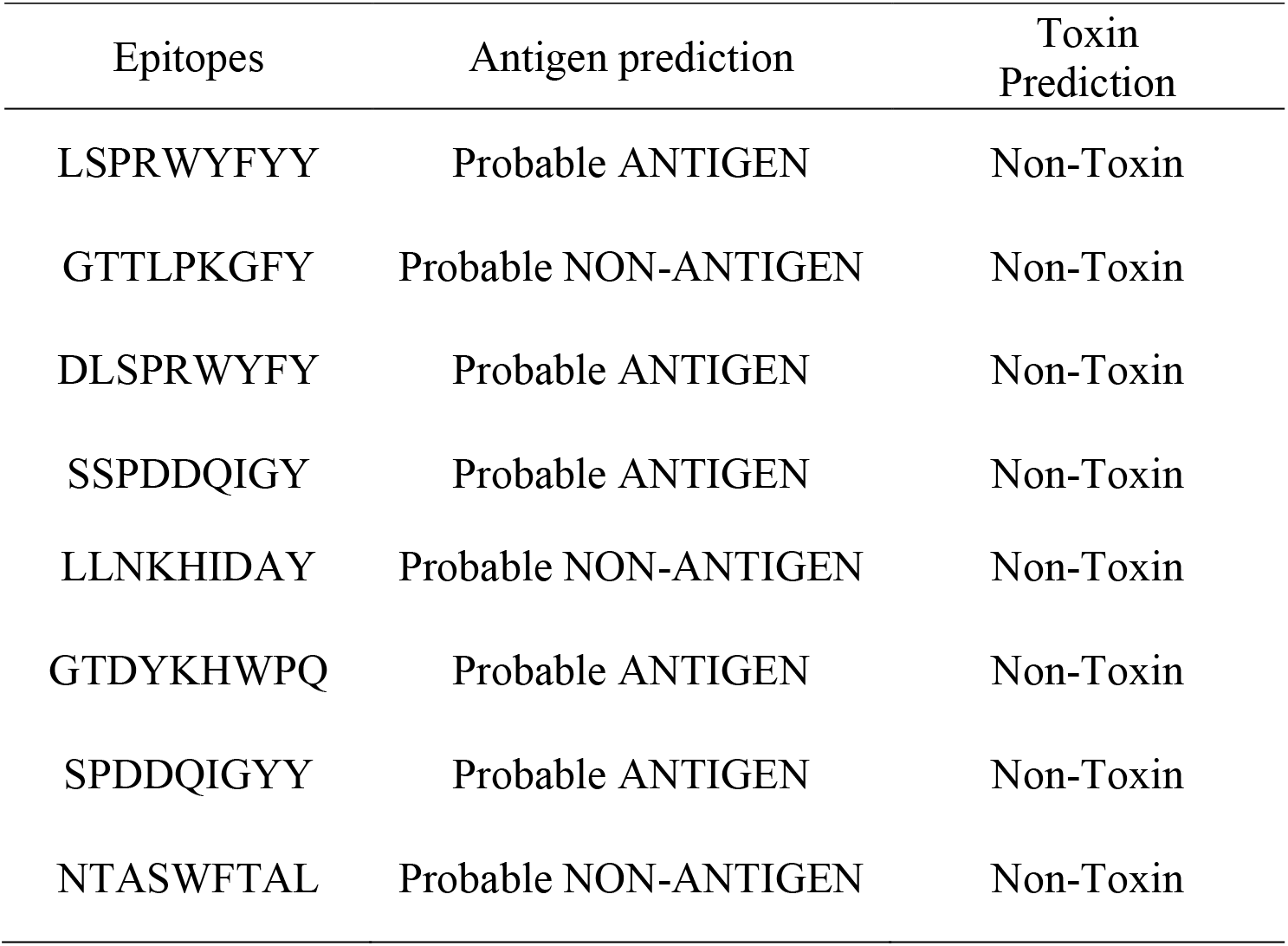
Antigenicity and toxicity prediction of selected epitopes

### 3.3 Toxicity prediction

The toxicity of the selected peptide sequences was assessed using the ToxinPred web server. The results from the ToxinPred server represented that all of our probable epitopes were found nontoxic **(Table 1)**.

### 3.4 Protein structure prediction and validation

The secondary of a protein describes the α-helix, β-sheets, and random coil. Our SARS-CoV-2 N protein has 419 residues, of which 213 residues (50.8%) from the helix, 187 residues from sheets (44.6%) and 66 (15.8%) residues from the coil **(Figure 3)**. For 3D structure, we built a model using EasyModeller, which was validated using PROCHECK and PROSA web server. The discrete optimized protein energy (DOPE) score was calculated −19585.62653. PROSA predicts the z-score of the model which was depicted 0.56 and the PROCHECK server was used for Ramachandran plot calculations. Ramachandran plot of the model protein indicated that 76% of residues in the most favorable region, 22% in the allowed region and 0.4% in the disallowed region **(Figure 4)**.

**FIGURE 3.**
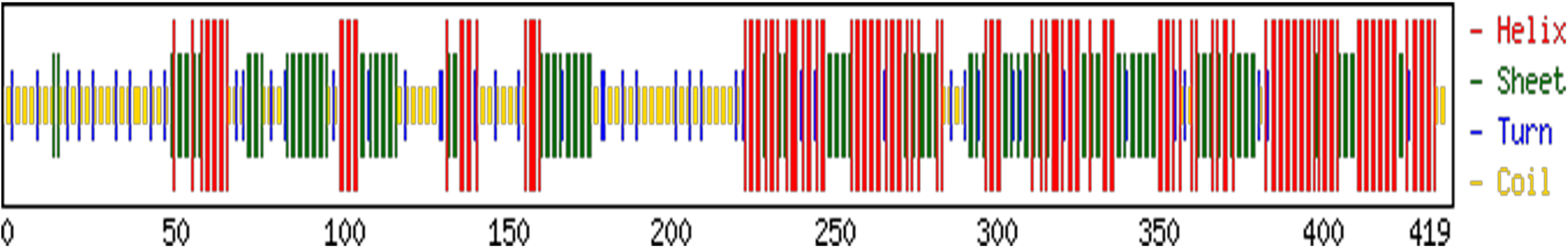
Composition of secondary structure from amino acid residues of SARS-CoV-2 nucleocapsid protein. Only 44.6% residues form sheet, 50.8% form helices and 15.8% residues form the turn region.

**FIGURE 4.**
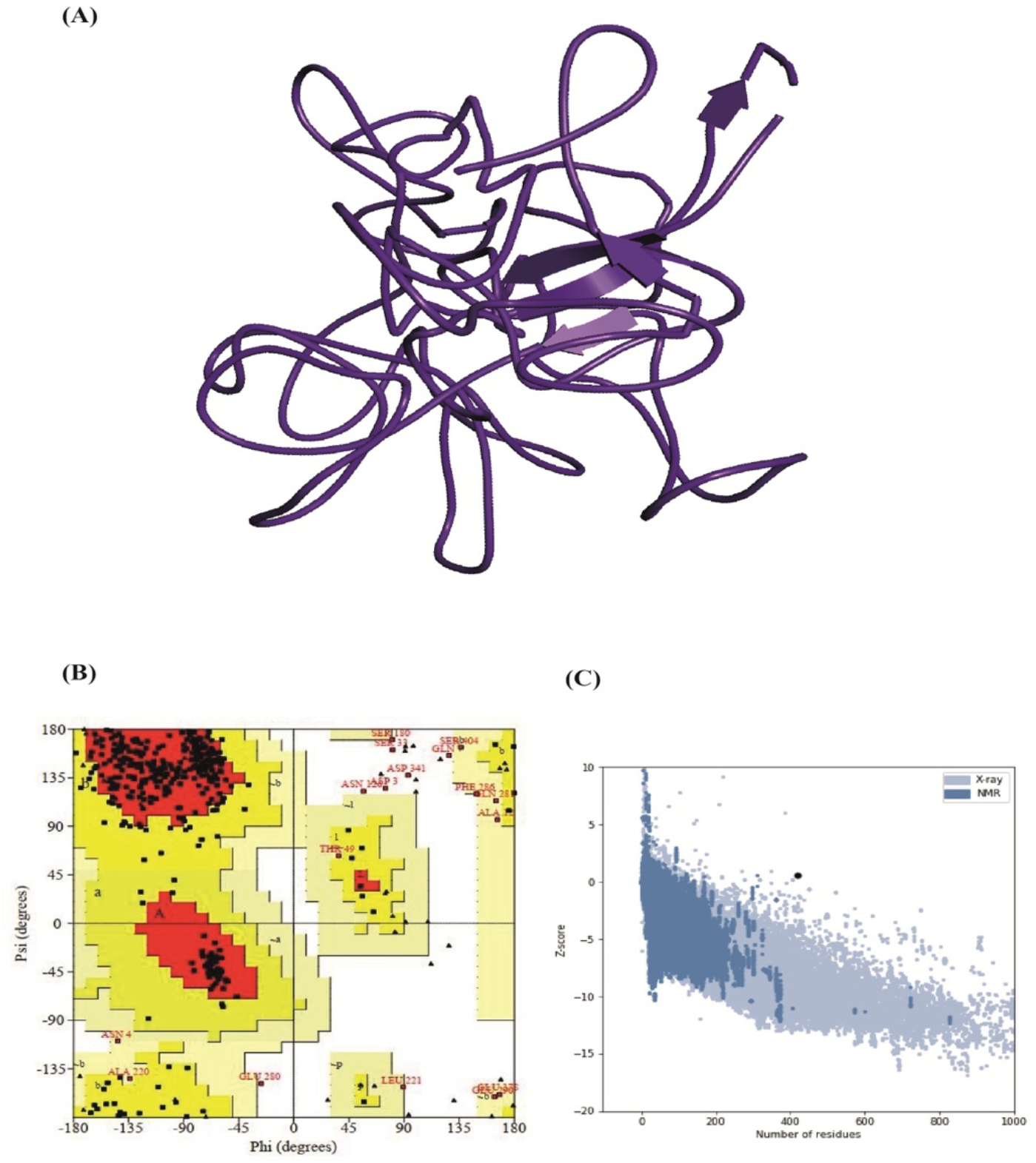
**(A)**The three-dimensional structure of the protein prepared using EasyModeller; **(B)** Ramachandran plot analysis of the protein using PROCHECK web server; **(C)** z-score predicted by PROSA server.

### 3.5 CD8^+^ T-cell epitope identification

Based on high combinatorial and MHC binding, the top eight epitopes were predicted by the NetCTL server from the selected protein sequence were selected for further analysis. Using the MHC-I binding prediction tool, which is based on SMM, we selected those MHC-I alleles for which the epitopes showed the highest affinity (IC_50_ < 200 nm). Proteasomes play an important role in cleaving the peptide bond, resulting in the conversion of protein into the peptide. The peptide molecules that are homogeneous to class I MHC molecules and the peptide-MHC molecule after the proteasomal cleavage were presented as T-helper cells after the transportation into the cell membrane. The total score of each epitope-HLA interaction was taken into consideration and higher processing efficiency was meant by obtaining a higher score. The epitope NTASWFTAL interacted with most of the MHC-I alleles including, HLA-A*68:02, HLA-C*16:01, HLA-C*03:03, HLA-C*03:04, HLA-C*12:03, HLA-A*02:06, HLA-C*03:02, HLA-A*26:01 and HLA-C*14:02 **(Table 2)**. Moreover, the MHC-NP prediction tool was used to find the highest probable score of our predicted epitope NTASWFTAL, with a score of 1.11, for HLA-A*68:02. Furthermore, all the predicted epitopes had a maximum identity for conservancy hit and 100% maximum identity was found **(Table 2)**. Also, the I-pMHC immunogenicity prediction analysis of the epitope NTASWFTAL was found 0.22775 **(Table 2)**.

**TABLE 2.**
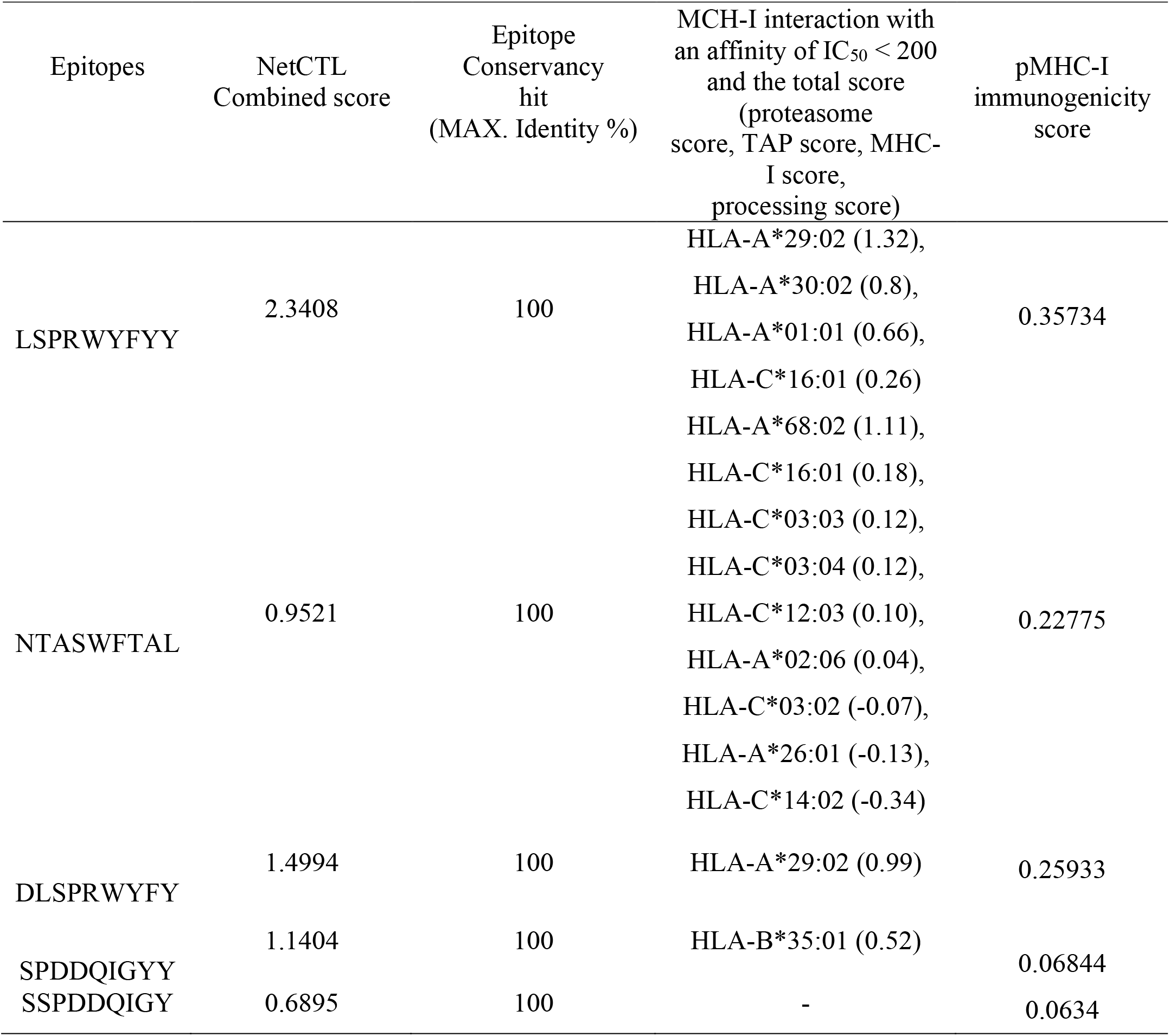
The potential CD8^+^ T-cell epitopes along with their interacting MHC class I alleles and total processing score, epitopes conservancy_hits and pMHC-I immunogenicity score

### 3.6 Population coverage

The cumulative amount of the population coverage was obtained for the predicted epitope NTASWFTAL Results from the population coverage demonstrated that with 57.16% coverage, East Asia found the highest coverage region. The results of the population coverage were shown in **Table 3 and Supplementary Figures S2-S5**.

**TABLE 3.** Analysis of the population coverage for the proposed epitope against SARS-Cov-2

| Population | Coverage(%)<sup>a</sup> | Average hit<sup>b</sup> | PC90<sup>c</sup> |
| --- | --- | --- | --- |
| Central Africa | 35.31 | 0.40 | 0.15 |
| East Africa | 39.25 | 0.45 | 0.16 |
| East Asia | 57.56 | 0.71 | 0.24 |
| Europe | 42.95 | 0.50 | 0.18 |
| North Africa | 42.15 | 0.49 | 0.17 |
| North America | 45.32 | 0.53 | 0.18 |
| Northeast Asia | 48.11 | 0.55 | 0.19 |
| Oceania | 31.43 | 0.34 | 0.15 |
| South Africa | 33.91 | 0.38 | 0.15 |
| South America | 38.66 | 0.44 | 0.16 |
| South Asia | 36.53 | 0.41 | 0.16 |
| Southeast Asia | 49.45 | 0.57 | 0.20 |
| Southwest Asia | 28.53 | 0.32 | 0.14 |
| West Africa | 56.22 | 0.67 | 0.23 |
| West Indies | 12.89 | 0.13 | 0.11 |

**Notes:** <sup>a</sup> Projected population coverage.
<sup>b</sup> Average number of epitope hits/HLA combinations recognized by the population.
<sup>c</sup> Minimum number of epitope hits/HLA combinations recognized by 90% of the population.

### 3.7 Allergenicity assessment

The AllerTop server was used for the identification of the allergic reaction caused by a vaccine in an individual which might be harmful or life-threatening. The allergenicity of the selected epitope was calculated using the AllerTop tool and predicted as probable non-allergen.

### 3.8 Molecular docking analysis for HLA and epitope interaction

In this study, the verification of the interaction between the HLA molecules and our predicted potential epitope was done by molecular docking simulation using Autodock Vina in PyRx 0.8 software. Among all the MHC class I alleles, only HLA-A*68:02 had a maximum probable score for our most potent epitope NTASWFTAL **(Figure 5)**. Therefore, we carried out the molecular docking study using HLA-A*68:02 (PDB ID: 4I48). We found that our predicted epitope NTASWFTAL interacted with HLA-A*68:02 with strong binding affinities of −9.4 kcal/mol **(Figure 6)**. The selected epitope interacted with Arg6, Ser4, Ser2, Asp30 residues of chain-A and Lys59, Asp60, Ser58, Gly30 of chain-B through hydrogen bonding (H-bond), whereas Lys7 residue of chain-B form bonds as a result of sharing electrons (which may happen as a result of charge distribution) **(Figure 6)**.

**FIGURE 5.**
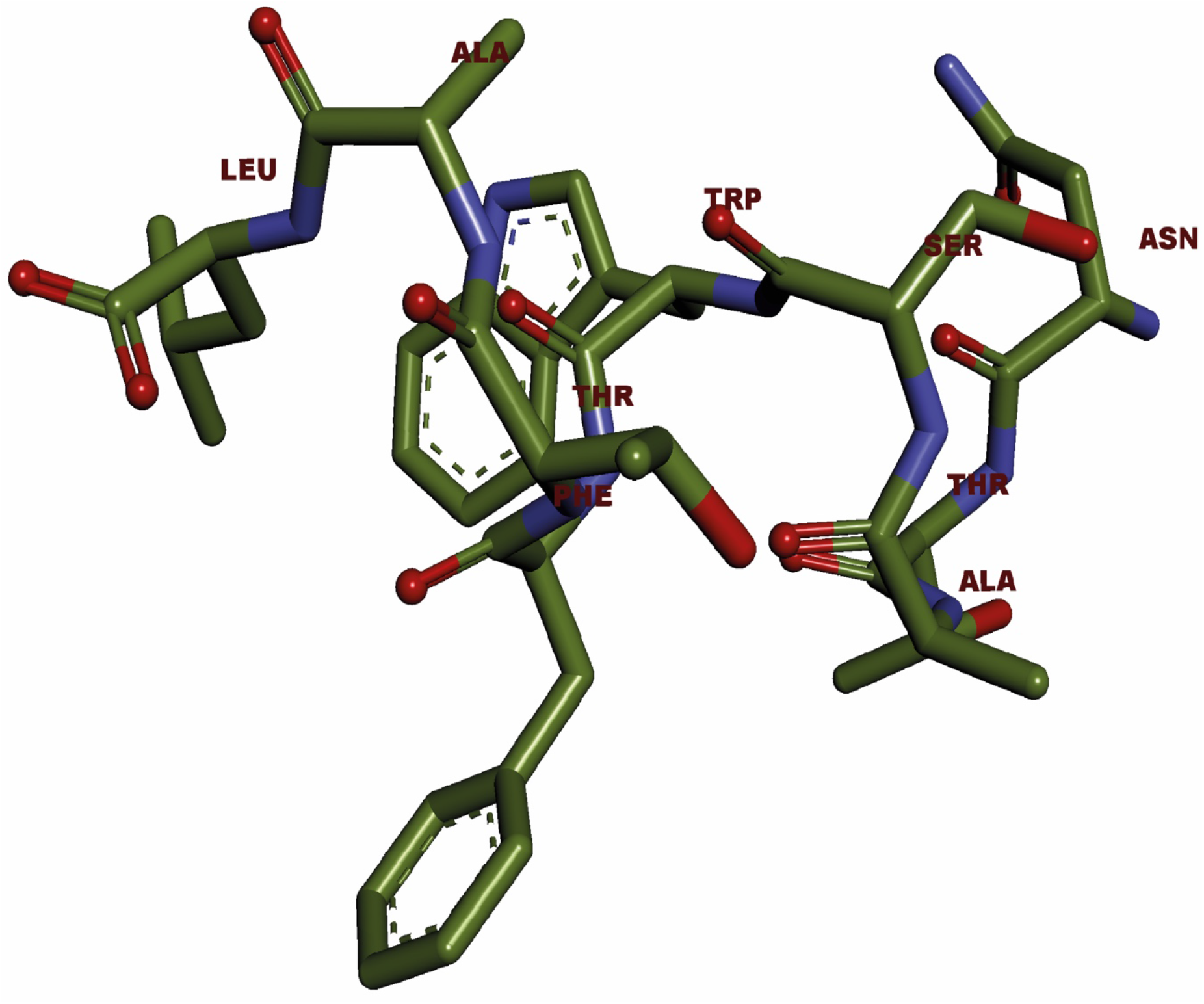
3D representation of our predicted epitope, NTASWFTAL.

**FIGURE 6.**
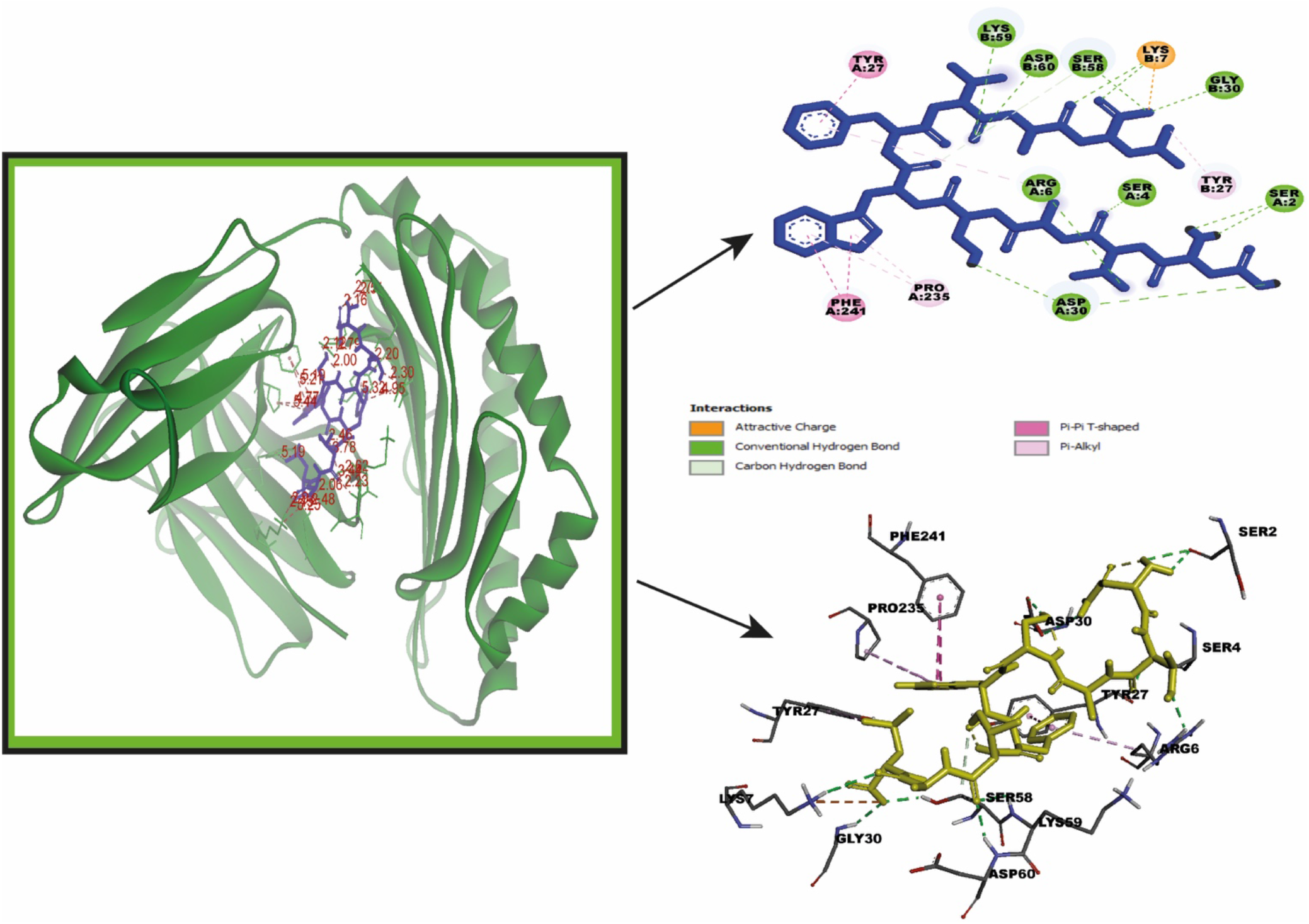
Molecular docking studies incorporating the binding of NTASWFTAL with groove of the HLA-A*68:02, both 2D and 3D representation. NTASWFTAL interacted with Arg6, Ser4, Ser2, Asp30 residues of chain-A and Lys59, Asp60, Ser58, Gly30 of chain-B through hydrogen bonding.

### 3.9 B-cell epitope prediction

In this study, using the amino acid scale-based method, we predicted the B-cell epitope identification. Different analysis methods were used for the prediction of continuous B-cell epitope. The results of the B-cell predictions were shown in **Table 4 and Figures 7-9**.

**FIGURE 7.**
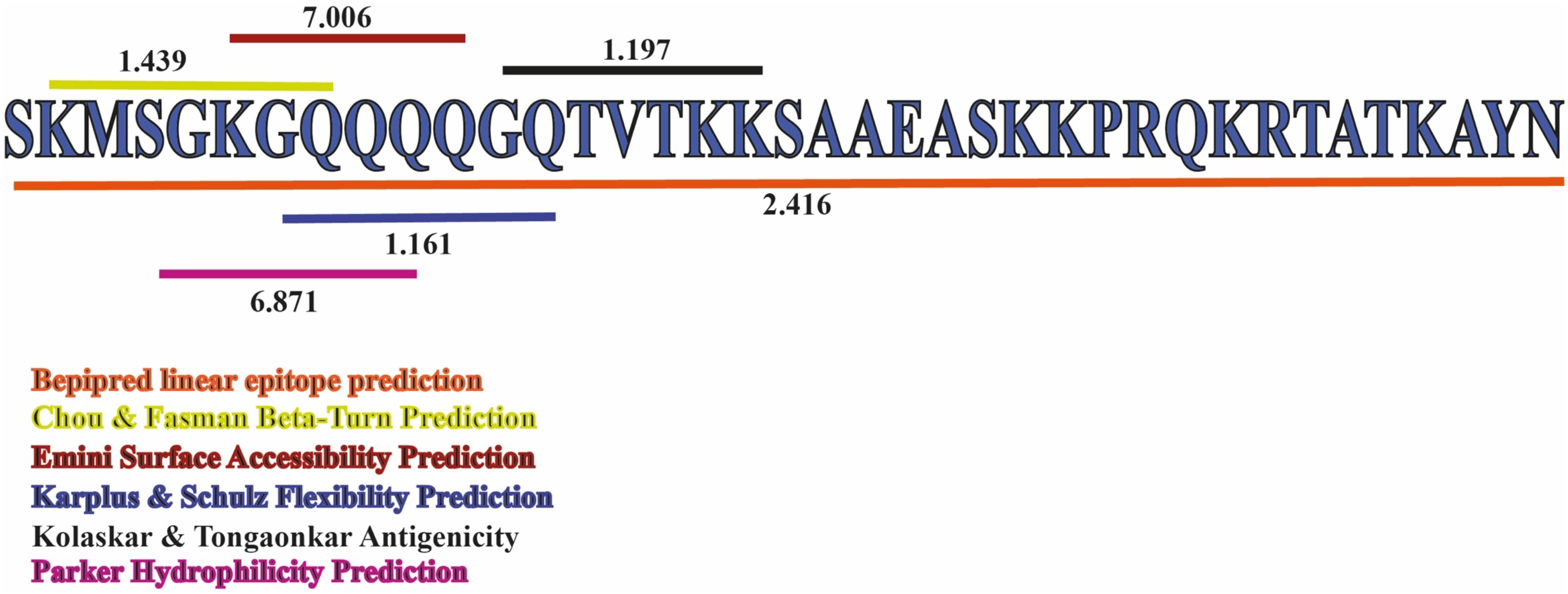
Combined B-cell linear epitope prediction showed the region from 232 to 249 amino acid residues had the highest antigenic propensity for B-cell linear epitopes. Surrounded by six differently coloured lines, which cover the region 232–269 amino acid residues in SARS-CoV-2 nucleocapsid pritein, each line indicating different analysis methods with the maximum scores.

**FIGURE 8.**
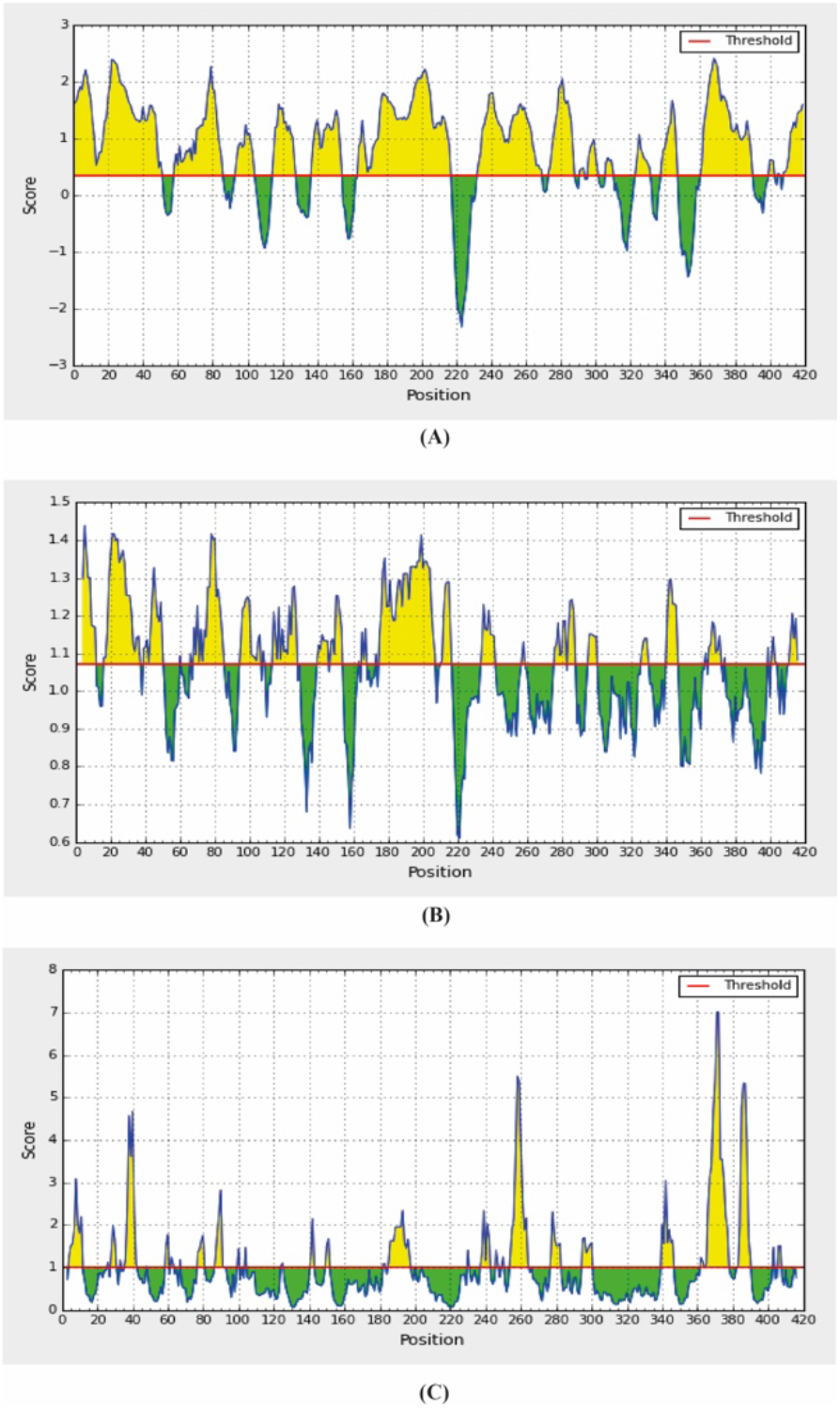
Combined B-cell linear epitope prediction using **(A)** Bepipred linear epitope prediction, **(B)** Chou & Fasman beta-turn prediction, **(C)** Emini surface accessibility prediction methods.

**FIGURE 9.**
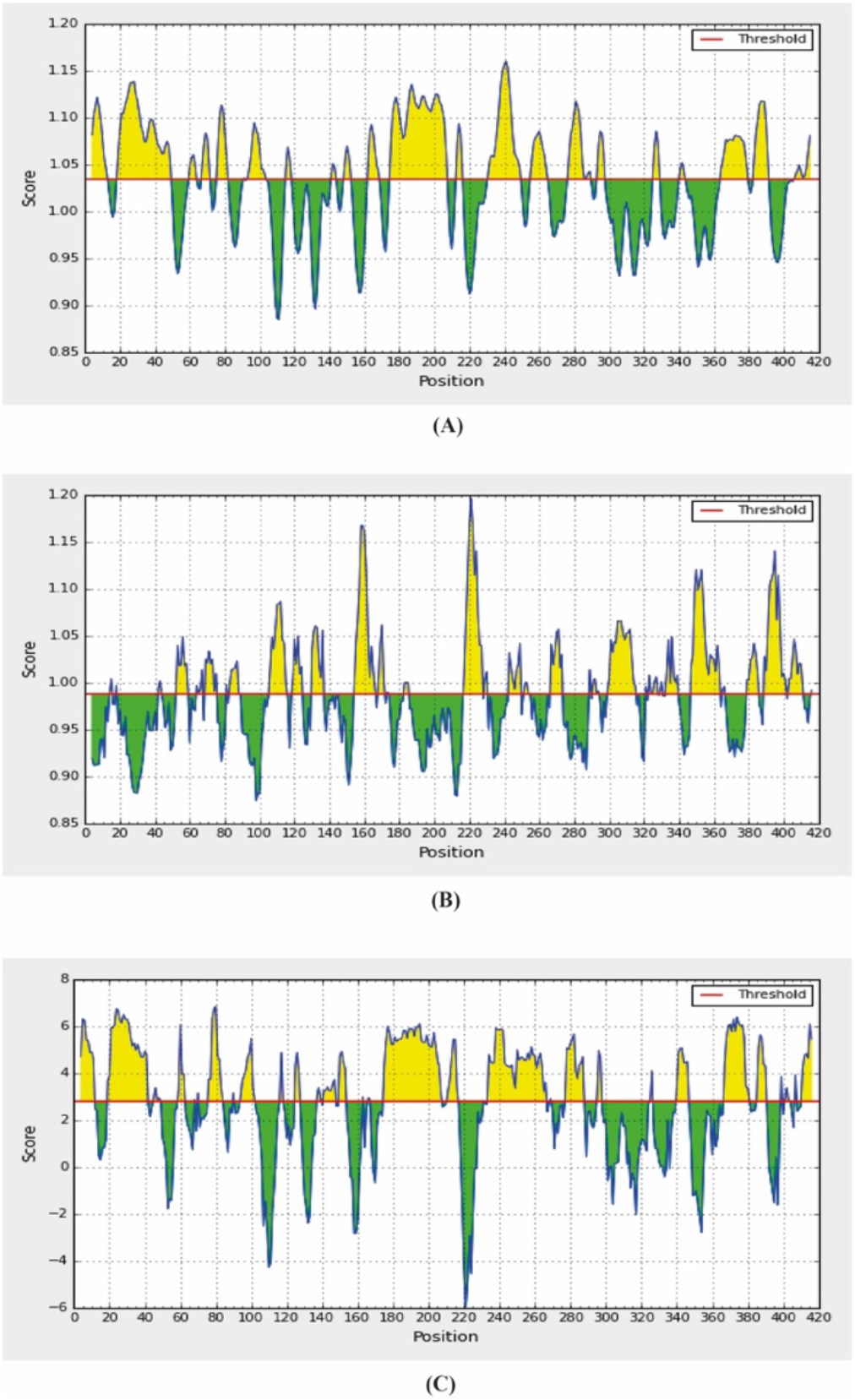
Combined B-cell linear epitope prediction using **(A)** Karplus & Schulz flexibility prediction, **(B)** Kolaskar & Tongaonkar antigenicity, **(C)** Parker hydrophilicity prediction methods.

**TABLE 4.** Combined B-cell linear epitope prediction

| Method | Region | Residues | Length | Score |  |  |
| --- | --- | --- | --- | --- | --- | --- |
|  |  |  |  | Max. | Avg. | Min. |
| Bepipred Linear Epitope Prediction | 232-269 | SKMSGKGQQQQ<br>GQTVTKKSAAEA<br>SKKPRQKRTATK<br>AYN | 38 | 2.416 | 0.813 | -0.001 |
| Chou & Fasman Beta-Turn Prediction | 233-239 | KMSGKGQ | 7 | 1.439 | 1.070 | 0.610 |
| Emini Surface Accessibility Prediction | 237-242 | KGQQQQ | 6 | 7.006 | 1.000 | 0.050 |
| Karplus & Schulz Flexibility Prediction | 238-244 | GQQQQGQ | 7 | 1.161 | 1.035 | 0.885 |
| Kolaskar & Tongaonkar Antigenicity | 243-249 | GQTVTKK | 7 | 1.197 | 0.988 | 0.874 |
| Parker Hydrophilicity Prediction | 235-241 | SGKGQQQ | 7 | 6.871 | 2.800 | -5.971 |

Firstly, Bepipred linear epitope prediction was used, which is regarded as the best single method for predicting linear B-cell epitopes using a Hidden Markov model. Our analysis revealed that the peptide sequences from 232 to 269 amino acid residues were able to induce the desired immune response as B-cell epitopes.

The β-turns were predicted by Chaus and Fasman β-turn prediction method. The region 233-239 residues were predicted as a β-turn region with a score of 1.164, which was higher than the average score.

For antigenicity prediction, the Kolaskar and Tongaonkar antigenicity prediction methods were implied. The method evaluates the antigenicity based on the physicochemical properties of amino acids and their abundances in experimentally known epitopes. The average antigenic propensity of our SARS-CoV-2 N protein was 0.988 with a maximum of 1.197 and a minimum of 0.874. Region 243-249 residues were found prone to antigenicity. Also, the average flexibility of 1.035 and a minimum of 0.874 was predicted by the Karplus and Schulz flexibility prediction method. The residues from 238 to 244 were found to be most flexible with the highest score of 1.161. The Parker Hydrophilicity Prediction tool predicts the hydrophilicity of SARS-CoV-2 N protein with an average score of 2.80, a minimum of 0.874 and the region 235-241 possessed a score of 5.943, which found moderately similar with the maximum value.

For predicting the surface ability, this study includes the Emini surface accessibility prediction method. The average surface accessibility was 1.0 and a minimum 0.050. In alignment with the previous B-cell epitope results, we predicted the peptide sequence from 237 to 242 had the better surface ability.

## 4 Discussion

As yet, it has been reported that the reproduction rate of SARS-CoV-2 is greater than SARS and MERS and the symptoms of the COVID-19 infection include fever with more than 38 °C body temperature along with alveolar edema, leading to difficulty in breathing, whereas mild symptoms perhaps not engender high fever (48). Surprisingly, with a high fatality rate, the severity of the infection was found more than the infection caused by both SARS and MERS, with multiple organ damage, which was reported not long ago (49).

At present, researchers are examining repurposed compounds from other viral infections to treat SARS-CoV-2. For example, both lopinavir and ritonavir are HIV protease inhibitors but in a lopinavir-ritonavir clinical trial report, the treatment benefit derived was dubious (50). From recovering patients, several convalescent immunoglobulins are derived which is currently investigated as a potential treatment for the disease (51). As there have been no approved treatments for COVID-19 that exists until now, these treatments are the best hope for striving to keep the mortality rate low before vaccines become widely available.

Despite many potential challenges, vaccine development is a crucial factor in modern biotechnology as vaccines are the most important prerequisites for defending the burden of diseases over the world (52). With the divulgement of sequence-based technology in genomics and proteomics, enough pieces of information are available regarding different eukaryotic and prokaryotic organisms including viruses. Therefore, utilizing various bioinformatics tools, it is possible to design peptide-based vaccines through comprehensibly studying the epitopes and several studies suggested epitope-based vaccines against different diseases including dengue, chikungunya, Saint Louis encephalitis virus (53–55). Although epitope-dependent vaccine design is quite familiar, a little research works are done in the case of SARS-CoV-2. Being an RNA virus, SARS-CoV-2 is different from the DNA virus as well as the rate of mutation is higher than the DNA viruses and according to various research, it can be assumed that the mutations might occur in the N protein (56). Recently, N proteins of SARS-CoV-2 are regarded as a primary target for vaccine development as its function includes viral replication and directly associated with the infection process (57), as a consequence related to the pathogenesis of COVID-19. Previous research works have already established that N proteins of several viruses as well as SARS are considered as a potential target for the development of vaccines (58–61). Moreover, we have already mentioned the detrimental role of SARS-CoV-2 in host-cell responses. This aspect leads us to conduct *in silico* experiments for designing a peptide-based vaccine against the novel SARS-CoV-2.

Earlier, it has been thought that vaccine development primarily relies on B-cell immunity, but recent discovery unveiled that T-cell epitopes are more propitious as a result of more long-lasting immune response mediated by CD8^+^ T-cells and due to the antigenic drift, by which an antibody is not able to respond against an antibody (62). In this study, focusing on MHC class I potential peptide epitopes, we predicted T-cell as well as B-cell epitopes which were able to show immune responses in various ways. Many characteristics including antigenicity, toxicity need to take into consideration for identifying a protein sequence-based epitope into a vaccine candidate and the predicted eight epitopes were fulfilled the entire criterion. However, only five potent epitopes have been predicted from the NetCTL 1.2 server and the epitopes were further taken for the progressive analysis. Besides, all peptides except SSPDDQIGY were able to interact with the MHC class I alleles, and NTASWFTAL interacted with the most MHC class I alleles. Amongst them, HLA-A*68:02 possessed the highest probable score. Further, the conservancy of the epitopes which was predicted by the IEDB conservancy analysis tool delineated that all of our predicted epitopes had the maximum identity of 100%. Therefore, we have taken the epitope NTASWFTAL for further analysis due to its maximum interaction with MHC class I alleles and the highest conservancy.

Allergenicity is regarded as one of the most noteworthy obstacles in vaccine development. Importantly, T-cells not CD4^+^ T-cells are involved in an allergic reaction and an allergic reaction is stimulated by type 2 T helper cell along with immunoglobulin E (63). In this experiment, we assessed the allergenicity using AllerTop 2.0, which is well recognized for its high sensitivity, and able to identify structurally diverse allergens in comparison with the known allergens. AllerTop predicted our selected epitope as non-allergen.

It has been proposed that the T-cell epitopes bind with the MHC molecules and MHC class I molecules generally presented short peptides that are 8-11 amino acid long, whereas MHC class II molecules present longer peptides with 13-17 amino acid residues (64). In this experiment, we determined the binding (presence of the antigen on the surface) affinity of the predicted epitope using molecular docking analysis and demonstrated that NTASWFTAL interacted with HLA-A*68:02 and found a binding affinity of −9.4 kcal/mol, which depicted a greater interaction with the epitope and the HLA molecule as the more negative energy implied to more binding affinity (65). The results from the molecular docking studies also revealed that epitope NTASWFTAL formed H-bond with both chain-A and chain-B of the HLA molecule and attractive charges were also responsible for the binding.

Another factor that is considered as the most prominent one during the process of vaccine development is population coverage, as the distribution of HLA varies according to ethnicity and geographical region. Our experiment showed that the epitope NTASWFTAL covered almost all regions of the world, where the highest coverage was observed in East Asia, where COVID-19 first reported. Interestingly, our findings indicated that our predicted epitope specifically binds with the widespread HLA molecules and the vaccine will be easily employed.

In addition, the B-cell epitope provides a strong immune response without causing any adverse effects. As a result, we also calculated the B-cell epitope prediction and found that the protein sequences from 232 to 249 amino acid residues as B-cell epitope. The identified region might be able to stimulate the desired immune response and also important for developing a vaccine.

## 5 Conclusions

The advancement in immunoinformatic has now emerged as a potential field for the prediction of epitope-based vaccines. As viruses can delineate both T-cell and humoral immunity, our predicted epitope might suggest enhancing the immunity against SARS-CoV-2. The assumption is based on the basic principles of immunity, which confers the attachment of virus with the host cell, evoking immune responses and transfers the information to a broad spectrum of T cells and B cells. Our investigated epitopes mimic the interaction to CD8 cells antigen presentation using computational approaches. However, our study is an introductory design to predict epitope-based vaccine against SARS-CoV-2 and we hope that this predicted epitope will assist the further laboratory analysis for designing and predicting novel candidates against COVID-19.

## 7 Author contributions

Study concept and design: A.R., S.A.S., M.A.I., and T.B.E. Acquisition of data: A.R., S.A.S., and M.A.I. Analyses and interpretation of data: A.R., M.A.I., and T.B.E. Drafting the manuscript: A.R., S.A.S., and T.B.E. Critical revision of the manuscript for important intellectual content: S.A., F.B.F., B.H.K., and T.B.E. Technical or material support: M.M.N.U. Study supervision: T.B.E.

## 9 Acknowledgments

The authors would like to thank Cambridge Proofreading^®^ & Editing LLC. (https://proofreading.org/) for editing a draft of this manuscript.

## 10 Data availability statement

The raw data supporting the conclusions of this manuscript will be made available by the authors, without undue reservation, to any qualified researcher.

## 13.1 Supplementary data

**Supplementary data 1** Multiple sequence alignment of SARS-CoV-2 nucleocapsid protein

## 13.2 Supplementary figures

**FIGURE S1** Evolutionary divergence analysis of available glycoproteins of different strains of SARS-CoV-2; results are represented in a phylogenetic tree.

**FIGURE S2** Population coverage based on MHC restriction data for **(A)** Central Africa, **(B)** East Africa, **(C)** East Asia, **(D)** North Africa - using the Immune Epitope Database analysis resource.

**FIGURE S3** Population coverage based on MHC restriction data for **(A)** North Africa, **(B)** North America, **(C)** Northeast Asia, **(D)** Oceania - using the Immune Epitope Database analysis resource.

**FIGURE S4** Population coverage based on MHC restriction data for **(A)** South Africa, **(B)** South America, **(C)** South Asia, **(D)** Southeast Asia - using the Immune Epitope Database analysis resource.

**FIGURE S5** Population coverage based on MHC restriction data for **(A)** Southwest Asia, **(B)** West Africa, **(C)** West Indies - using the Immune Epitope Database analysis resource.

## REFERENCES

1. Wang D, Hu B, Hu C, Zhu F, Liu X, Zhang J, et al. Clinical characteristics of 138 hospitalized patients with 2019 novel coronavirus--infected pneumonia in Wuhan, China. Jama. 2020. doi:10.1001/jama.2020.1585

2. Li Q, Guan X, Wu P, Wang X, Zhou L, Tong Y, et al. Early transmission dynamics in Wuhan, China, of novel coronavirus--infected pneumonia. New England Journal of Medicine. 2020. doi: 10.1056/NEJMoa2001316

3. Gralinski LE, Menachery VD. Return of the Coronavirus: 2019-nCoV. Viruses. 2020;12(2):135. doi: 10.3390/v12020135

4. Organization WH, others. Coronavirus disease 2019 (COVID-19): situation report, 72. 2020;

5. Zhu H, Guo Q, Li M, Wang C, Fang Z, Wang P, et al. Host and infectivity prediction of Wuhan 2019 novel coronavirus using deep learning algorithm. BioRxiv. 2020. doi: 10.1101/2020.01.21.914044

6. Wu F, Zhao S, Yu B, Chen Y-M, Wang W, Song Z-G, et al. A new coronavirus associated with human respiratory disease in China. Nature. 2020;579(7798):265–9. doi: 10.1038/s41586-020-2008-3

7. Huang C, Wang Y, Li X, Ren L, Zhao J, Hu Y, et al. Clinical features of patients infected with 2019 novel coronavirus in Wuhan, China. The Lancet. 2020. doi: 10.1016/S0140-6736(20)30183-5

8. Chan JF-W, Yuan S, Kok K-H, To KK-W, Chu H, Yang J, et al. A familial cluster of pneumonia associated with the 2019 novel coronavirus indicating person-to-person transmission: a study of a family cluster. The Lancet. 2020;395(10223):514–23. doi: 10.1016/S0140-6736(20)30154-9

9. Chan JF-W, Kok K-H, Zhu Z, Chu H, To KK-W, Yuan S, et al. Genomic characterization of the 2019 novel human-pathogenic coronavirus isolated from a patient with atypical pneumonia after visiting Wuhan. Emerging microbes & infections. 2020;9(1):221–36. doi: 10.1080/22221751.2020.1719902

10. Wu F, Zhao S, Yu B, Chen Y-M, Wang W, Hu Y, et al. Complete genome characterisation of a novel coronavirus associated with severe human respiratory disease in Wuhan, China. bioRxiv. 2020. doi: 10.1101/2020.01.24.919183

11. Narayanan K, Huang C, Makino S. SARS coronavirus accessory proteins. Virus research. 2008;133(1):113–21. doi: 10.1016/j.virusres.2007.10.009

12. Dermime S, Gilham DE, Shaw DM, Davidson EJ, Meziane E-K, Armstrong A, et al. Vaccine and antibody-directed T cell tumour immunotherapy. Biochimica et Biophysica Acta (BBA)-Reviews on Cancer. 2004;1704(1):11–35. doi: 10.1016/j.bbcan.2004.03.002

13. Meloen RH, Langeveld JPM, Schaaper WMM, Slootstra JW. Synthetic peptide vaccines: unexpected fulfillment of discarded hope? Biologicals. 2001;29(3–4):233–6. doi: 10.1006/biol.2001.0298

14. Channappanavar R, Fett C, Zhao J, Meyerholz DK, Perlman S. Virus-specific memory CD8 T cells provide substantial protection from lethal severe acute respiratory syndrome coronavirus infection. Journal of virology. 2014;88(19):11034–44. doi: 10.1128/JVI.01505-14

15. Rappuoli R, Black S, Bloom DE. Vaccines and global health: In search of a sustainable model for vaccine development and delivery. Science Translational Medicine. 2019;11(497):eaaw2888. doi: 10.1126/scitranslmed.aaw2888

16. Olsson S-E, Villa LL, Costa RLR, Petta CA, Andrade RP, Malm C, et al. Induction of immune memory following administration of a prophylactic quadrivalent human papillomavirus (HPV) types 6/11/16/18 L1 virus-like particle (VLP) vaccine. Vaccine. 2007;25(26):4931–9. doi: 10.1016/j.vaccine.2007.03.049

17. Suarez DL, Schultz-Cherry S. Immunology of avian influenza virus: a review. Developmental & Comparative Immunology. 2000;24(2–3):269–83. doi: 10.1016/S0145-305X(99)00078-6

18. Briney B, Sok D, Jardine JG, Kulp DW, Skog P, Menis S, et al. Tailored immunogens direct affinity maturation toward HIV neutralizing antibodies. Cell. 2016;166(6):1459–70. doi: 10.1016/j.cell.2016.08.005

19. Pedersen SR, Christensen JP, Buus S, Rasmussen M, Korsholm KS, Nielsen M, et al. Immunogenicity of HLA class I and II double restricted influenza A-derived peptides. PloS one. 2016;11(1). doi: 10.1371/journal.pone.0145629

20. Ying LIN, Xu S, Yang RF, Li YX, Ji YY, He YY, et al. Identification of an epitope of SARS-coronavirus nucleocapsid protein. Cell research. 2003;13(3):141–5. doi: 10.1038/sj.cr.7290158

21. Johnson M, Zaretskaya I, Raytselis Y, Merezhuk Y, McGinnis S, Madden TL. NCBI BLAST: a better web interface. Nucleic acids research. 2008;36(suppl_2):W5--W9. doi: 10.1093/nar/gkn201

22. Li W, Cowley A, Uludag M, Gur T, McWilliam H, Squizzato S, et al. The EMBL-EBI bioinformatics web and programmatic tools framework. Nucleic acids research. 2015;43(W1):W580--W584. doi: 10.1093/nar/gkv279

23. Tamura K, Stecher G, Peterson D, Filipski A, Kumar S. MEGA6: molecular evolutionary genetics analysis version 6.0. Molecular biology and evolution. 2013;30(12):2725–9. doi: 10.1093/molbev/mst197

24. Doytchinova IA, Flower DR. VaxiJen: a server for prediction of protective antigens, tumour antigens and subunit vaccines. BMC bioinformatics. 2007;8(1):4. doi: 10.1186/1471-2105-8-4

25. Gupta S, Kapoor P, Chaudhary K, Gautam A, Kumar R, Raghava GPS. Peptide toxicity prediction. In: Computational Peptidology. Springer; 2015. p. 143–57.

26. Kumar TA. CFSSP: Chou and Fasman secondary structure prediction server. Wide Spectrum. 2013;1(9):15–9. doi: 10.1007/978-1-4939-2285-7_7

27. Van Brussel H, Wyns J, Valckenaers P, Bongaerts L, Peeters P. Reference architecture for holonic manufacturing systems: PROSA. Computers in industry. 1998;37(3):255–74. doi: 10.1016/S0166-3615(98)00102-X

28. Kuntal BK, Aparoy P, Reddanna P. EasyModeller: A graphical interface to MODELLER. BMC Research Notes. 2010; doi: 10.1186/1756-0500-3-226

29. Webb B, Sali A. Comparative protein structure modeling using MODELLER. Current protocols in bioinformatics. 2016;54(1):5–6. doi: 10.1002/cpbi.3

30. Laskowski RA, MacArthur MW, Moss DS, Thornton JM. PROCHECK: a program to check the stereochemical quality of protein structures. Journal of Applied Crystallography. 1993; doi: 10.1107/S0021889892009944

31. Buus S, Lauemøller SL, Worning P, Kesmir C, Frimurer T, Corbet S, et al. Sensitive quantitative predictions of peptide-MHC binding by a ‘Query by Committee’artifìcial neural network approach. Tissue antigens. 2003;62(5):378–84.

32. Giguère S, Drouin A, Lacoste A, Marchand M, Corbeil J, Laviolette F. MHC-NP: predicting peptides naturally processed by the MHC. Journal of immunological methods. 2013;400:30–6. doi: 10.1034/j.1399-0039.2003.00112.x

33. Bui H-H, Sidney J, Li W, Fusseder N, Sette A. Development of an epitope conservancy analysis tool to facilitate the design of epitope-based diagnostics and vaccines. BMC bioinformatics. 2007;8(1):361. doi: 10.1186/1471-2105-8-361

34. Moutaftsi M, Peters B, Pasquetto V, Tscharke DC, Sidney J, Bui H-H, et al. A consensus epitope prediction approach identifies the breadth of murine T CD8+-cell responses to vaccinia virus. Nature biotechnology. 2006;24(7):817–9. doi: 10.1038/nbt1215

35. Dimitrov I, Flower DR, Doytchinova I. AllerTOP-a server for in silico prediction of allergens. In: BMC bioinformatics. 2013. p. S4. doi: 10.1186/1471-2105-14-S6-S4

36. Maupetit J, Derreumaux P, Tuffery P. PEP-FOLD: an online resource for de novo peptide structure prediction. Nucleic acids research. 2009;37(suppl_2):W498--W503. doi: 10.1093/nar/gkp323

37. Dallakyan S. PyRx-python prescription v. 0.8. The Scripps Research Institute. 2008;2010.

38. Dunbrack RL. Rotamer libraries in the 21st century. Current Opinion in Structural Biology. 2002. doi: 10.1016/S0959-440X(02)00344-5

39. Pettersen EF, Goddard TD, Huang CC, Couch GS, Greenblatt DM, Meng EC, et al. UCSF Chimera - A visualization system for exploratory research and analysis. Journal of Computational Chemistry. 2004; doi: 10.1002/jcc.20084

40. O’Boyle NM, Banck M, James CA, Morley C, Vandermeersch T, Hutchison GR. Open Babel: An open chemical toolbox. Journal of cheminformatics. 2011;3(1):33. doi: 10.1186/1758-2946-3-33

41. Trott O, Olson AJ. AutoDock Vina: Improving the speed and accuracy of docking with a new scoring function, efficient optimization, and multithreading. Journal of Computational Chemistry. 2009; doi: 10.1002/jcc.21334

42. Chou PY, Fasman GD. Empirical predictions of protein conformation. Annual review of biochemistry. 1978;47(1):251–76.

43. Emini EA, Hughes J V, Perlow D, Boger J. Induction of hepatitis A virus-neutralizing antibody by a virus-specific synthetic peptide. Journal of virology. 1985;55(3):836–9.

44. Kolaskar AS, Tongaonkar PC. A semi-empirical method for prediction of antigenic determinants on protein antigens. FEBS letters. 1990;276(1–2):172–4. doi: 10.1016/0014-5793(90)80535-Q

45. Karplus PA, Schulz GE. Prediction of chain flexibility in proteins. Naturwissenschaften. 1985;72(4):212–3. doi: 10.1007/BF01195768

46. Larsen JEP, Lund O, Nielsen M. Improved method for predicting linear B-cell epitopes. Immunome research. 2006;2(1):2. doi: 10.1186/1745-7580-2-2

47. Rini JM, Schulze-Gahmen U, Wilson IA. Structural evidence for induced fit as a mechanism for antibody-antigen recognition. Science. 1992;255(5047):959–65. doi: 10.1126/science.1546293

48. Liu Y, Gayle AA, Wilder-Smith A, Rocklöv J. The reproductive number of COVID-19 is higher compared to SARS coronavirus. Journal of travel medicine. 2020. doi: 10.1093/jtm/taaa021

49. Wang T, Du Z, Zhu F, Cao Z, An Y, Gao Y, et al. Comorbidities and multi-organ injuries in the treatment of COVID-19. The Lancet. 2020. doi: 10.1016/S0140-6736(20)30558-4

50. Cao B, Wang Y, Wen D, Liu W, Wang J, Fan G, et al. A trial of lopinavir--ritonavir in adults hospitalized with severe Covid-19. New England Journal of Medicine. 2020. doi: 10.1056/NEJMoa2001282

51. Chen L, Xiong J, Bao L, Shi Y. Convalescent plasma as a potential therapy for COVID-19. The Lancet Infectious Diseases. 2020;20(4):398–400. doi: 10.1016/S1473-3099(20)30141-9

52. Oany AR, Emran A-A, Jyoti TP. Design of an epitope-based peptide vaccine against spike protein of human coronavirus: an in silico approach. Drug design, development and therapy. 2014;8:1139. doi: 10.2147/DDDT.S67861

53. Hasan A, Hossain M, Alam J. A computational assay to design an epitope-based Peptide vaccine against Saint Louis encephalitis virus. Bioinformatics and Biology insights. 2013;7:BBI--S13402. doi: 10.4137/BBI.S13402

54. Chakraborty S, Chakravorty R, Ahmed M, Rahman A, Waise TM, Hassan F, et al. A computational approach for identification of epitopes in dengue virus envelope protein: a step towards designing a universal dengue vaccine targeting endemic regions. In silico biology. 2010;10(5, 6):235–46. doi: 10.3233/ISB-2010-0435

55. Islam R, Sakib MS, Zaman A. A computational assay to design an epitope-based peptide vaccine against chikungunya virus. Future Virology. 2012;7(10):1029–42. doi: 10.2217/fvl.12.95

56. Huang Y, Khorchid A, Wang J, Parniak MA, Darlix J-L, Wainberg MA, et al. Effect of mutations in the nucleocapsid protein (NCp7) upon Pr160 (gag-pol) and tRNA (Lys) incorporation into human immunodeficiency virus type 1. Journal of virology. 1997;71(6):4378–84.

57. Thomas JA, Gorelick RJ. Nucleocapsid protein function in early infection processes. Virus research. 2008;134(1–2):39–63. doi: 10.1016/j.virusres.2007.12.006

58. Kim TW, Lee JH, Hung C-F, Peng S, Roden R, Wang M-C, et al. Generation and characterization of DNA vaccines targeting the nucleocapsid protein of severe acute respiratory syndrome coronavirus. Journal of virology. 2004;78(9):4638–45. doi: 10.1128/JVI.78.9.4638-4645.2004

59. Sabara M, Frenchick PJ, Mullin-Ready KF. Rotavirus nucleocapsid protein VP6 in vaccine compositions. Biotechnology Advances. 1995;13(4):803–4.

60. Arthur LO, Bess Jr JW, Chertova EN, Rossio JL, Esser MT, Benveniste RE, et al. Chemical inactivation of retroviral infectivity by targeting nucleocapsid protein zinc fingers: a candidate SIV vaccine. AIDS research and human retroviruses. 1998;14:S311--9.

61. Zhao P, Cao J, Zhao L-J, Qin Z-L, Ke J-S, Pan W, et al. Immune responses against SARS-coronavirus nucleocapsid protein induced by DNA vaccine. Virology. 2005;331(1):128–35. doi: 10.1016/j.virol.2004.10.016

62. Chiou S-S, Fan Y-C, Crill WD, Chang R-Y, Chang G-JJ. Mutation analysis of the cross-reactive epitopes of Japanese encephalitis virus envelope glycoprotein. Journal of general virology. 2012;93(6):1185–92. doi: 10.1099/vir.0.040238-0

63. Kallinich T, Beier KC, Wahn U, Stock P, Hamelmann E. T-cell co-stimulatory molecules: their role in allergic immune reactions. European Respiratory Journal. 2007;29(6):1246–55. doi: 10.1183/09031936.00094306

64. Alberts B, Johnson A, Lewis J, Raff M, Roberts K, Peter Walter P. Molecular Biology of the Cell, New York: Garland Science.[Google Scholar]. 2002;

65. Ahmed S, Rakib A, Islam MA, Khanam BH, Faiz FB, Paul A, et al. In vivo and in vitro pharmacological activities of Tacca integrifolia rhizome and investigation of possible lead compounds against breast cancer through in silico approaches. Clinical Phytoscience [Internet]. 2019;5(1):36. doi: 10.1186/s40816-019-0127-x

