## Supplemental Materials for "Epitope-Based Peptide Vaccine Against Severe Acute Respiratory Syndrome-Coronavirus-2 Nucleocapsid Protein: An *in silico* Approach"

### *Supplementary Materials*

#### Table of Contents

**Supplementary data**

**Supplementary data 1** Multiple sequence alignment of SARS-CoV-2 nucleocapsid protein

**Supplementary figures**

**FIGURE S1** Evolutionary divergence analysis of available glycoproteins of different strains of SARS-CoV-2; results are represented in a phylogenetic tree.

**FIGURE S2** Population coverage based on MHC restriction data for **(A)** Central Africa, **(B)** East Africa, **(C)** East Asia, **(D)** North Africa - using the Immune Epitope Database analysis resource.

**FIGURE S3** Population coverage based on MHC restriction data for **(A)** North Africa, **(B)** North America, **(C)** Northeast Asia, **(D)** Oceania - using the Immune Epitope Database analysis resource.

**FIGURE S4** Population coverage based on MHC restriction data for **(A)** South Africa, **(B)** South America, **(C)** South Asia, **(D)** Southeast Asia - using the Immune Epitope Database analysis resource.

**FIGURE S5** Population coverage based on MHC restriction data for **(A)** Southwest Asia, **(B)** West Africa, **(C)** West Indies - using the Immune Epitope Database analysis resource

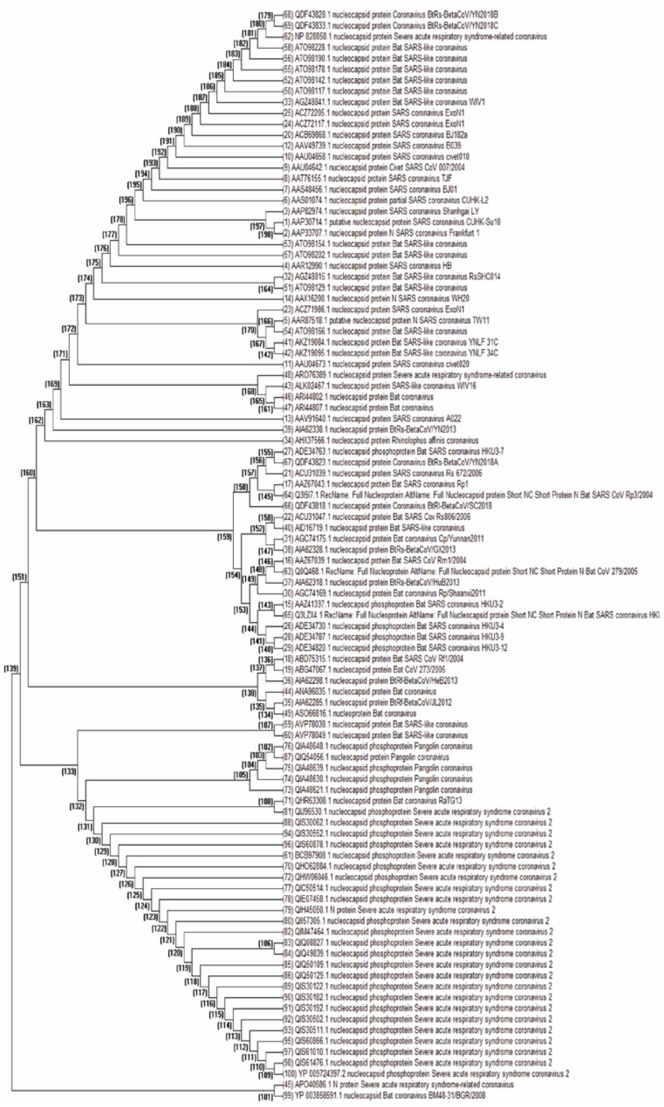

**FIGURE S1** Evolutionary divergence analysis of available glycoproteins of different strains of SARS-CoV-2; results are represented in a phylogenetic tree.

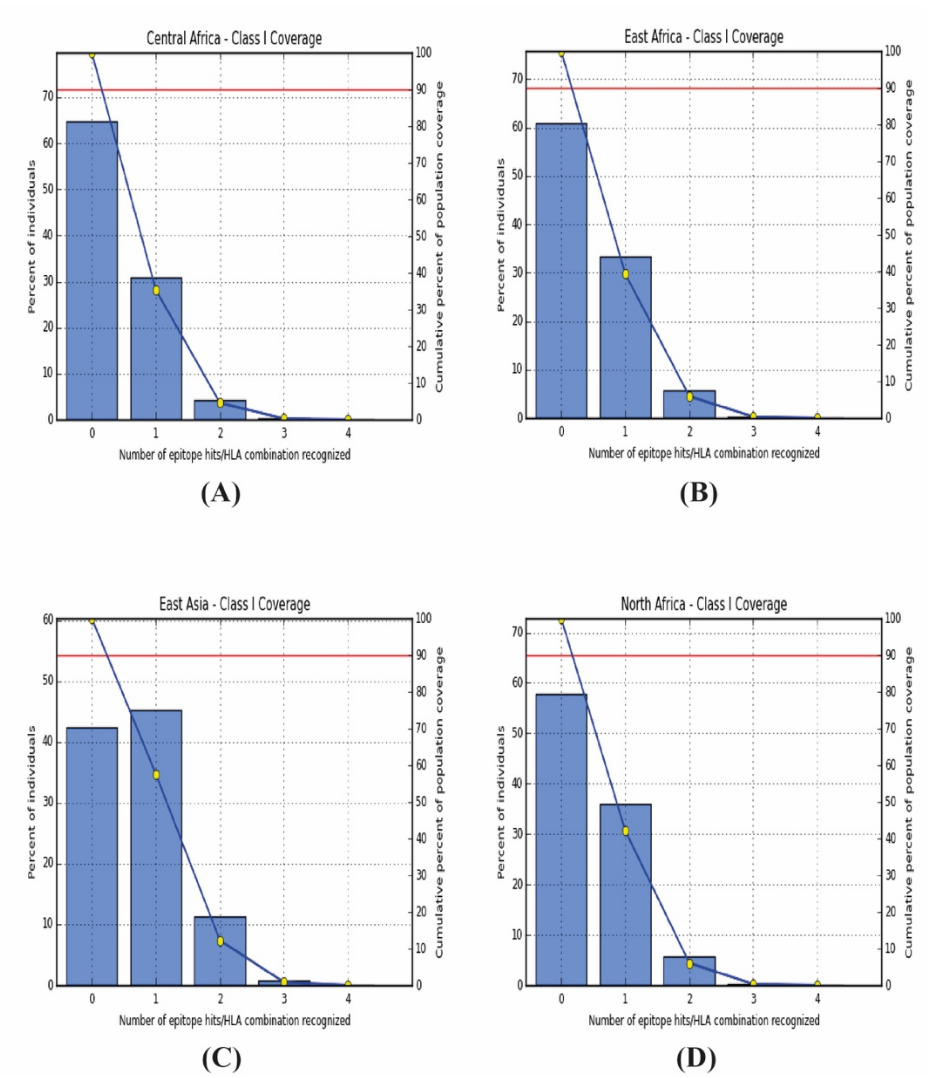


**FIGURE S2** Population coverage based on MHC restriction data for **(A)** Central Africa, **(B)** East Africa, **(C)** East Asia, **(D)** North Africa - using the Immune Epitope Database analysis resource.

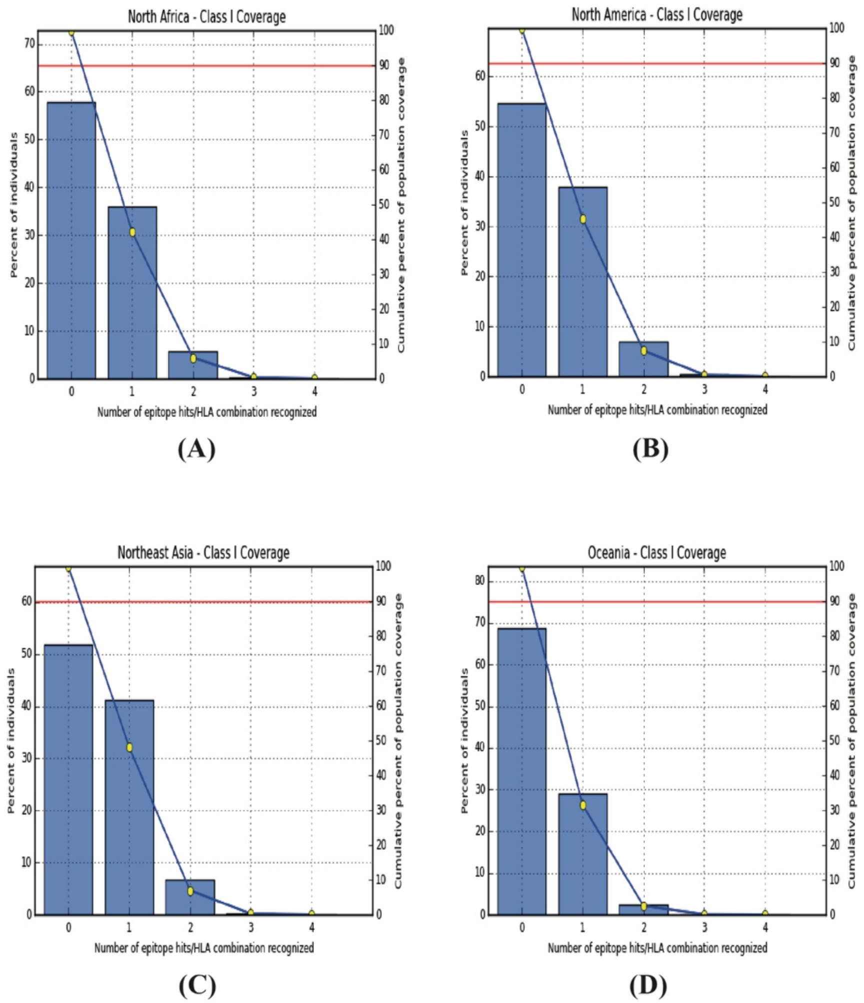


**FIGURE S3** Population coverage based on MHC restriction data for **(A)** North Africa, **(B)** North America, **(C)** Northeast Asia, **(D)** Oceania - using the Immune Epitope Database analysis resource.

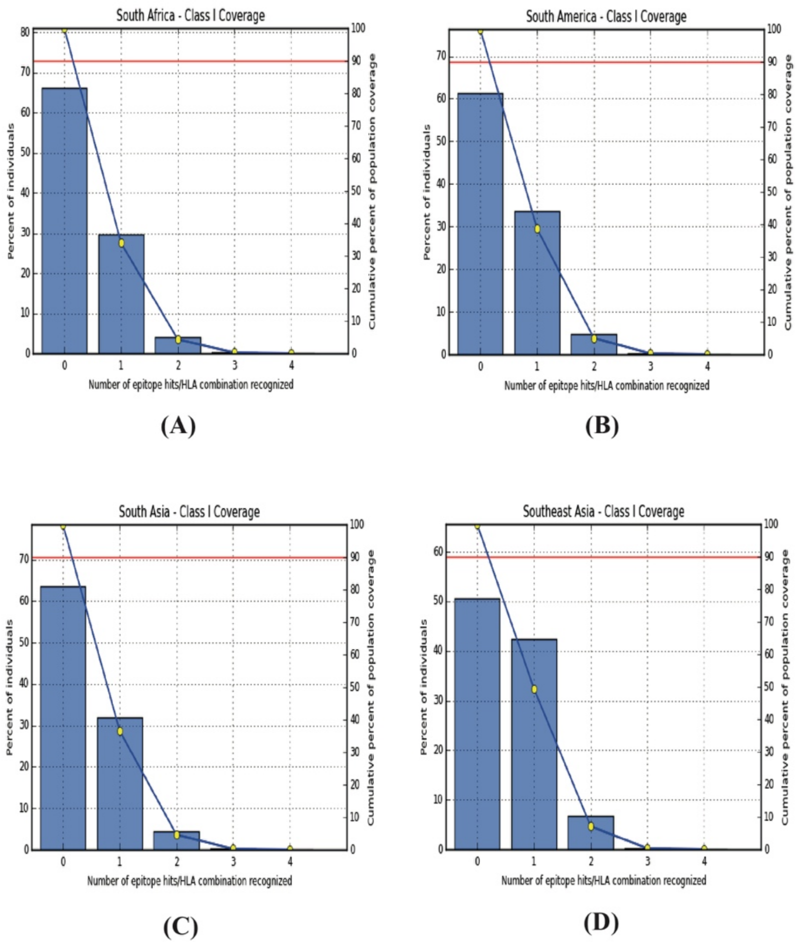


**FIGURE S4** Population coverage based on MHC restriction data for **(A)** South Africa, **(B)** South America, **(C)** South Asia, **(D)** Southeast Asia - using the Immune Epitope Database analysis resource.

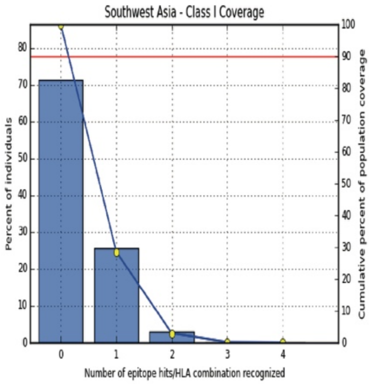


**(A)**

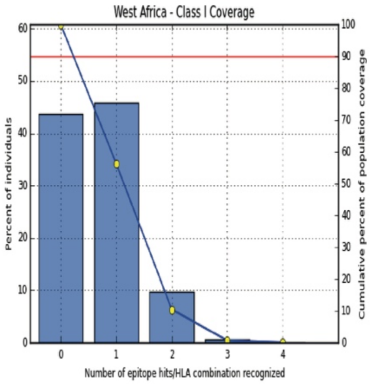


**(B)**

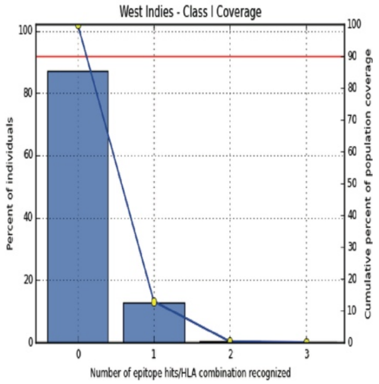


**(C)**

**FIGURE S5** Population coverage based on MHC restriction data for **(A)** Southwest Asia, **(B)** West Africa, **(C)** West Indies - using the Immune Epitope Database analysis resource.

### *Multiple Sequence Alignment*

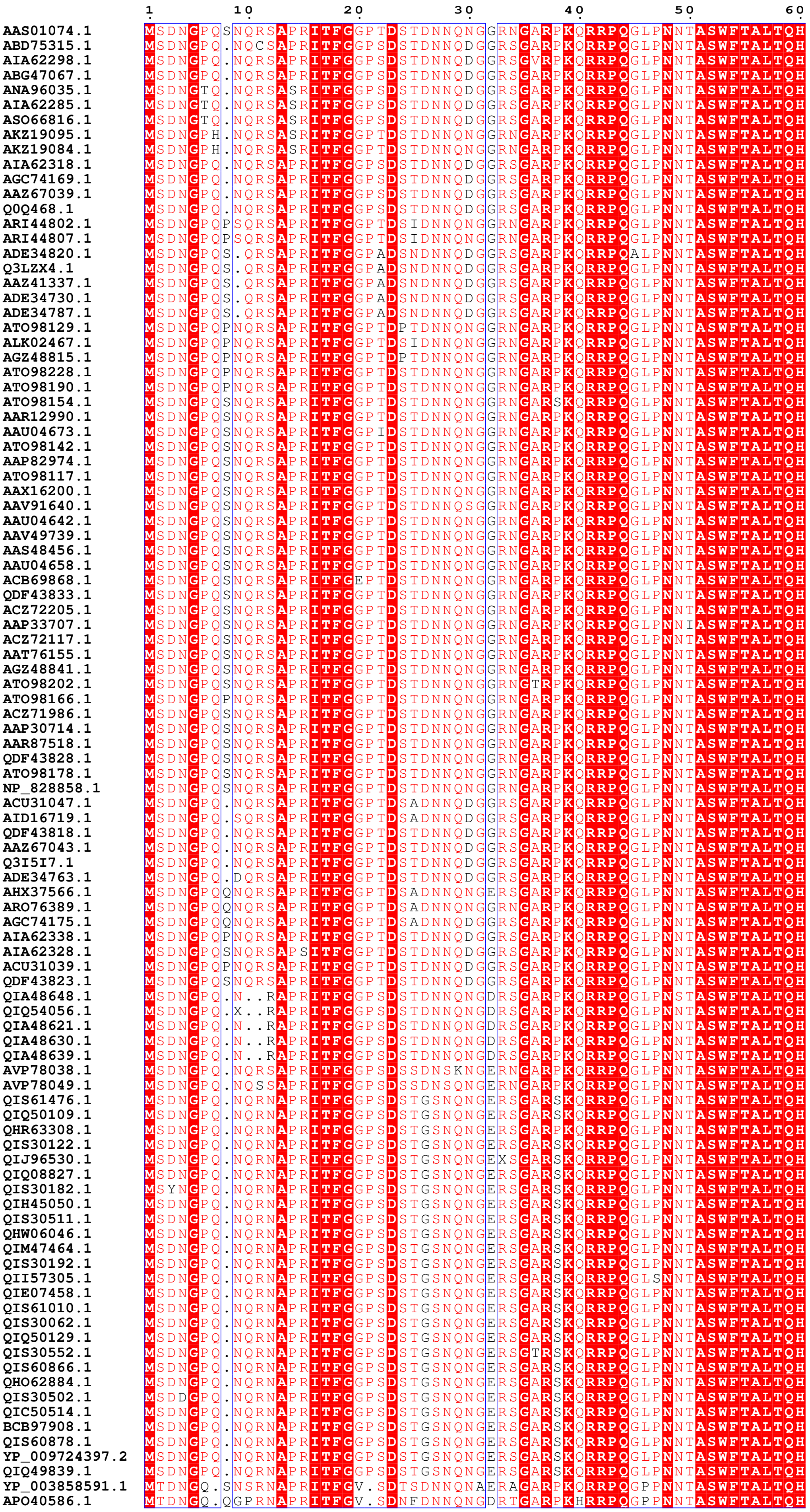

```
                          70        80        90       100       110       120
AAS01074.1        GKEELRFPRGQGVPINTNSGPDDQIGYYRRATRRVRGGDGKMKELSPRWYFYYLGTGPEA
ABD75315.1        GKEGLKFPQGQGVPINTNSGRDDQIGYYRRATRRVRGGDGKMKELSPRWYFYYLGTGPEA
AIA62298.1        GKEGLKFPQGQGVPINTNSGRDDQIGYYRRATRRVRGGDGKMKELSPRWYFYYLGTGPEA
ABG47067.1        GKEGLKFPQGQGVPINTNSGRDDQIGYYRRATRRVRGGDGKMKELSPRWYFYYLGTGPEA
ANA96035.1        GKEGLKFPQGQGVPINTNSGTDDQIGYYRRATRRVRGGDGKMKELSPRWYFYYLGTGPEA
AIA62285.1        GKEGLKFPQGQGVPINTNSGRDDQIGYYRRATRRVRGGDGKMKELSPRWYFYYLGTGPEA
ASO66816.1        GKEGLKFPQGQGVPINTNSGRDDQIGYYRRATRRVRGGDGKMKELSPRWYFYYLGTGPEA
AKZ19095.1        GKEELRFPQGQGVPINTNSGPDDQIGYYRRATRRVRGGDGKMKELSPRWYFYYLGTGPEA
AKZ19084.1        GKEELRFPQGQGVPINTNSGPDDQIGYYRRATRRVRGGDGKMKELSPRWYFYYLGTGPEA
AIA62318.1        GKEELRFPRGQGVPINTNSGKDDQIGYYRRATRRVRGGDGKMKELSPRWYFYYLGTGPEA
AGC74169.1        GKEELRFPRGQGVPINTNSGKDDQIGYYRRATRRVRGGDGKMKELSPRWYFYYLGTGPEA
AAZ67039.1        GKEELRFPRGQGVPINTNSGKDDQIGYYRRATRRVRGGDGKMKELSPRWYFYYLGTGPEA
Q0Q468.1          GKEELRFPRGQGVPINTNSGKDDQIGYYRRATRRVRGGDGKMKKLSPRWYFYYLGTGPEA
ARI44802.1        GKEELRFSRGQGVPINTNSGPDDQIGYYRRATRRVRGGDGKMKELSPRWYFYYLGTGPEA
ARI44807.1        GKEELRFSRGQGVPINTNSGPDDQIGYYRRATRRVRGGDGKMKELSPRWYFYYLGTGPEA
ADE34820.1        GKEELRFPRGQGVPINTNSGKDDQIGYYRRATRRVRGGDGKMKELSPRWYFYYLGTGPEA
Q3LZX4.1          GKEELRFPRGQGVPINTNSGKDDQIGYYRRATRRVRGGDGKMKELSPRWYFYYLGTGPEA
AAZ41337.1        GKEELRFPRGQGVPINTNSGKDDQIGYYRRATRRVRGGDGKMKELSPRWYFYYLGTGPEA
ADE34730.1        GKEELRFPRGQGVPINTNSGKDDQIGYYRRATRRVRGGDGKMKELSPRWYFYYLGTGPEA
ADE34787.1        GKEELRFPRGQGVPINTNSGKDDQIGYYRRATRRVRGGDGKMKELSPRWYFYYLGTGPEA
ATO98129.1        GKEELRFPRGQGVPINTNSGPDDQIGYYRRATRRVRGGDGKMKELSPRWYFYYLGTGPEA
ALK02467.1        GKEELRFPRGQGVPINTNSGPDDQIGYYRRATRRVRGGDGKMKELSPRWYFYYLGTGPEA
AGZ48815.1        GKEELRFPRGQGVPINTNSGPDDQIGYYRRATRRVRGGDGKMKELSPRWYFYYLGTGPEA
ATO98228.1        GKEELRFPRGQGVPINTNSGPDDQIGYYRRATRRVRGGDGKMKELSPRWYFYYLGTGPEA
ATO98190.1        GKEELRFPRGQGVPINTNSGPDDQIGYYRRATRRVRGGDGKMKELSPRWYFYYLGTGPEA
ATO98154.1        GKEELRFPRGQGVPINTNSGPDDQIGYYRRATRRVRGGDGKMKELSPRWYFYYLGTGPEA
AAR12990.1        GKEELRFPRGQGVPINTNSGPDDQIGYYRRATRRARGGDGKMKELSPRWYFYYLGTGPEA
AAU04673.1        GKEELRFPRGQGVPINTNSGPDDQIGYYRRATRRVRGGDGKMKELSPRWYFYYLGTGPEA
ATO98142.1        GKEELRFPRGQGVPINTNSGPDDQIGYYRRATRRVRGGDGKMKELSPRWYFYYLGTGPEA
AAP82974.1        GKEELRFPRGQGVPINTNSGPDDQIGYYRRATRRVRGGDGKMKELSPRWYFYYLGTGPEA
ATO98117.1        GKEELRFPRGQGVPINTNSGPDDQIGYYRRATRRVRGGDGKMKELSPRWYFYYLGTGPEA
AAX16200.1        GKEELRFPRGQGVPINTNSGPDDQIGYYRRATRRVRGGDGKMKELSPRWYFYYLGTGPEA
AAV91640.1        GKEELRFPRGQGVPINTNSGPDDQIGYYRRATRRVRGGDGKMKELSPRWYFYYLGTGPEA
AAU04642.1        GKEELRFPRGQGVPINTNSGPDDQIGYYRRATRRVRGGDGKMKELSPRWYFYYLGTGPEA
AAV49739.1        GKEELRFPRGQGVPINTNSGPDDQIGYYRRATRRVRGGDGKMKELSPRWYFYYLGTGPEA
AAS48456.1        GKEELRFPRGQGVPINTNSGPDDQIGYYRRATRRVRGGDGKMKELSPRWYFYYLGTGPEA
AAU04658.1        GKEELRFPRGQGVPINTNSGPDDQIGYYRRATRRVRGGDGKMKELSPRWYFYYLGTGPEA
ACB69868.1        GKEELRFPRGQGVPINTNSGPDDQIGYYRRATRRVRGGDGKMKELSPRWYFYYLGTGPEA
QDF43833.1        GKEELRFPRGQGVPINTNSGPDDQIGYYRRATRRVRGGDGKMKELSPRWYFYYLGTGPEA
ACZ72205.1        GKEELRFPRGQGVPINTNSGPDDQIGYYRRATRRVRGGDGKMKELSPRWYFYYLGTGPEA
AAP33707.1        GKEELRFPRGQGVPINTNSGPDDQIGYYRRATRRVRGGDGKMKELSPRWYFYYLGTGPEA
ACZ72117.1        GKEELRFPRGQGVPINTNSGPDDQIGYYRRATRRVRGGDGKMKELSPRWYFYYLGTGPEA
AAT76155.1        GKEELRFPRGQGVPINTNSGPDDQIGFYRRATRRVRGGDGKMKELSPRWYFYYLGTGPEA
AGZ48841.1        GKEELRFPRGQGVPINTNSGPDDQIGYYRRATRRVRGGDGKMKELSPRWYFYYLGTGPEA
ATO98202.1        GKEELRFPRGQGVPINTNSGPDDQIGYYRRATRRVRGGDGKMKELSPRWYFYYLGTGPEA
ATO98166.1        GKEELRFPRGQGVPINTNSGPDDQIGYYRRATRRVRGGDGKMKELSPRWYFYYLGTGPEA
ACZ71986.1        GKEELRFPRGQGVPINTNSGPDDQIGYYRRATRRVRGGDGKMKELSPRWYFYYLGTGPEA
AAP30714.1        GKEELRFPRGQGVPINTNSGPDDQIGYYRRATRRVRGGDGKMKELSPRWYFYYLGTGPEA
AAR87518.1        GKEELRFPRGQGVPINTNSGPDDQIGYYRRATRRVRGGDGKMKELSPRWYFYYLGTGPEA
QDF43828.1        GKEELRFPRGQGVPINTNSGPDDQIGYYRRATRRVRGGDGKMKELSPRWYFYYLGTGPEA
ATO98178.1        GKEELRFPRGQGVPINTNSGPDDQIGYYRRATRRVRGGDGKMKELSPRWYFYYLGTGPEA
NP_828858.1       GKEELRFPRGQGVPINTNSGPDDQIGYYRRATRRVRGGDGKMKELSPRWYFYYLGTGPEA
ACU31047.1        GKEELRFPRGQGVPINTNSGKDDQIGYYRRATRRVRGGDGKMKELSPRWYFYYLGTGPEA
AID16719.1        GKEELRFPRGQGVPINTNSGKDDQIGYYRKATRRVRGGDGKMKELSPRWYFYYLGTGPEA
QDF43818.1        GKEELKFPRGQGVPINTNSGKDDQIGYYRRATRRVRGGDGKMKELSPRWYFYYLGTGPEA
AAZ67043.1        GKEELRFPRGQGVPINTNSGKDDQIGYYRRATRRVRGGDGKMKELSPRWYFYYLGTGPEA
Q3I5I7.1          GKEELRFPRGQGVPINTNSGKDDQIGYYRRATRRVRGGDGKMKELSPRWYFYYLGTGPEA
ADE34763.1        GKEELRFPRGQGVPINTNSGKDDQIGYYRRATRRVRGGDGKMKELSPRWYFYYLGTGPEA
AHX37566.1        GKEELRFPRGQGVPINTNSGTDDQIGYYRRATRRVRGGDGKMKELSPRWYFYYLGTGPEA
ARO76389.1        GKEELRFPRGQGVPINTNSGPDDQIGYYRRATRRVRGGDGKMKELSPRWYFYYLGTGPEA
AGC74175.1        GKEELRFPRGQGVPINTNSGKDDQIGYYRRATRRVRGGDGKMKELSPRWYFYYLGTGPEA
AIA62338.1        GKEELRFPRGQGVPINTNSGPDDQIGYYRRATRRVRGGDGKMKELSPRWYFYYLGTGPEA
AIA62328.1        GKEELRFPRGQGVPINTNSGKDDQIGYYRRATRRVRGGDGKMKELSPRWYFYYLGTGPEA
ACU31039.1        GKEELRFPRGQGVPINTNSGKDDQIGYYRRATRRVRGGDGKMKELSPRWYFYYLGTGPEA
QDF43823.1        GKEELRFPRGQGVPINTNSGKDDQIGYYRRATRRVRGGDGKMKELSPRWYFYYLGTGPEA
QIA48648.1        GKEDLRFPRGQGVPINTNSTKDDQIGYYRRATRRVRGGDGKMKDLSPRWYFYYLGTGPEA
QIQ54056.1        GKEDLRFPRGQGVPINTNSTKDDQIGYYRRATRRVRGGDGKMKDLSPRWYFYYLGTGPEA
QIA48621.1        GKEDLRFPRGQGVPINTNSTKDDQIGYYRRATRRVRGGDGKMKDLSPRWYFYYLGTGPEA
QIA48630.1        GKEDLRFPRGQGVPINTNSTKDDQIGYYRRATRRVRGGDGKMKDLSPRWYFYYLGTGPEA
QIA48639.1        GKEDLRFPRGQGVPINTNSTKDDQIGYYRRATRRVRGGDGKMKDLSPRWYFYYLGTGPEA
AVP78038.1        GKENLTFPRGQGVPINTNSSKDDQIGYYRRATRRIRGGDGKMKELSPRWYFYYLGTGPEA
AVP78049.1        GKENLTFPRGQGVPINTNSSKDDQIGYYRRATRRIRGGDGKMKELSPRWYFYYLGTGPEA
QIS61476.1        GKEDLKFPRGQGVPINTNSSPDDQIGYYRRATRRIRGGDGKMKDLSPRWYFYYLGTGPEA
QIQ50109.1        GKEDLKFPRGQGVPINTNSSPDXXXXYYXXATRRIRGGDGKMKDLSPRWYFYYLGTGPEA
QHR63308.1        GKEDLKFPRGQGVPINTNSSPDDQIGYYRRATRRIRGGDGKMKDLSPRWYFYYLGTGPEA
QIS30122.1        GKEDLKFPRGQGVPINTNSSPDDQIGYYRRATRRIRGGDGKMKDLSPRWYFYYLGTGPEA
QIJ96530.1        GKEDLKFPRGQGVPINTNSSPDDQIGYYRRATRRIRGGDGKMKDLSPRWYFYYLGTGPEA
QIQ08827.1        GKEDLKFPRGQGVPINTNSSPDDQIGYYRRATRRIRGGDGKMKDLSPRWYFYYLGTGPEA
QIS30182.1        GKEDLKFPRGQGVPINTNSSPDDQIGYYRRATRRIRGGDGKMKDLSPRWYFYYLGTGPEA
QIH45050.1        GKEDLKFPRGQGVPINTNSSPDDQIGYYRRATRRIRGGDGKMKDLSPRWYFYYLGTGPEA
QIS30511.1        GKEDLKFPRGQGVPINTNSSPDDQIGYYRRATRRIRGGDGKMKDLSPRWYFYYLGTGPEA
QHW06046.1        GKEDLKFPRGQGVPINTNSSPDDQIGYYRRATRRIRGGDGKMKDLSPRWYFYYLGTGPEA
QIM47464.1        GKEDLKFPRGQGVPINTNSSPDDQIGYYRRATRRIRGGDGKMKDLSPRWYFYYLGTGPEA
QIS30192.1        GKEDLKFPRGQGVPINTNSSPDDQIGYYRRATRRIRGGDGKMKDLSPRWYFYYLGTGPEA
QII57305.1        GKEDLKFPRGQGVPINTNSSPDDQIGYYRRATRRIRGGDGKMKDLSPRWYFYYLGTGPEA
QIE07458.1        GKEDLKFPRGQGVPINTNSSPDDQIGYYRRATRRIRGGDGKMKDLSPRWYFYYLGTGPEA
QIS61010.1        GKEDLKFPRGQGVPINTNSSPDDQIGYYRRATRRIRGGDGKMKDLSPRWYFYYLGTGPEA
QIS30062.1        GKEDLKFPRGQGVPINTNSSPDDQIGYYRRATRRIRGGDGKMKDLSPRWYFYYLGTGPEA
QIQ50129.1        GKEDLKFPRGQGVPINTNSSPDDQIGYYRRATRRIRGGDGKMKDLSPRWYFYYLGTGPEA
QIS30552.1        GKEDLKFPRGQGVPINTNSSPDDQIGYYRRATRRIRGGDGKMKDLSPRWYFYYLGTGPEA
QIS60866.1        GKEDLKFPRGQGVPINTNSSPDDQIGYYRRATRRIRGGDGKMKDLSPRWYFYYLGTGPEA
QHO62884.1        GKEDLKFPRGQGVPINTNSSPDDQIGYYRRATRRIRGGDGKMKDLSPRWYFYYLGTGPEA
QIS30502.1        GKEDLKFPRGQGVPINTNSSPDDQIGYYRRATRRIRGGDGKMKDLSPRWYFYYLGTGPEA
QIC50514.1        GKEDLKFPRGQGVPINTNSSPDDQIGYYRRATRRIRGGDGKMKDLSPRWYFYYLGTGPEA
BCB97908.1        GKEDLKFPRGQGVPINTNSSPDDQIGYYRRATRRIRGGDGKMKDLSPRWYFYYLGTGPEA
QIS60878.1        GKEDLKFPRGQGVPINTNSSPDDQIGYYRRATRRIRGGDGKMKDLSPRWYFYYLGTGPEA
YP_009724397.2    GKEDLKFPRGQGVPINTNSSPDDQIGYYRRATRRIRGGDGKMKDLSPRWYFYYLGTGPEA
QIQ49839.1        GKEDLKFPRGQGVPINTNSSPDDQIGYYRRATRRIRGGDGKMKDLSPRWYFYYLGTGPEA
YP_003858591.1    GKEGLSFPRGQGVPVNTNSTRDDQIGYYRRATRRVRGGDGKMKELSPRWYFYYLGTGPEA
APO40586.1        GKETLTFPRGQGVPINTNSGKDDQIGYYRRASRRVRGGDGKMKELSPRWYFYYLGTGPEA
```

```
                       130       140       150       160       170       180
AAS01074.1      SLPYGANKEGIVWVATEXXXXXXXXXXXXXXXXXXXXXXXXLPQGTTLPKGFYAEGSRGG
ABD75315.1      SLPYGANKEGIVWVATEGALNTPKDHIGTRNPNNNAAIVLQLPQGTTLPKGFYAEGSRNG
AIA62298.1      SLPYGANKEGIVWVATEGALNTPKDHIGTRNPNNNAAIVLQLPQGTTLPKGFYAEGSRNG
ABG47067.1      SLPYGANKEGIVWVATEGALNTPKDHIGTRNPNNNAAIVLQLPQGTTLPKGFYAEGSRNG
ANA96035.1      SLPYGANKEGIVWVATEGALNTPKDHIGTRNPNNNAAIVLQLPQGTTLPKGFYAEGSRSG
AIA62285.1      SLPYGANKEGIVWVATEGALNTPKDHIGTRNPNNNAAIVLQLPQGTTLPKGFYAEGSRSG
ASO66816.1      SLPYGANKEGIVWVAIEGALNTPKDHIGTRNPNNNAAIVLQLPQGTTLPKGFYAEGSRSG
AKZ19095.1      SLPYGANKEGIIWVATEGALNTPKDHIGTRNPNNNAATVLQLPQGTTLPKGFYAEGSRGG
AKZ19084.1      SLPYGANKEGIIWVATEGALNTPKDHIGTRNPNNNAATVLQLPQGTTLPKGFYAEGSRGG
AIA62318.1      SLPYGANKEGIVWVATEGALNTPKDHIGTRNPNNNAAIVLQLPQGTTLPKGFYAEGSRGG
AGC74169.1      SLPYGANKEGIIWVATEGALNTPKDHIGTRNPNNNAAIVLQLPQGTTLPKGFYAEGSRGG
AAZ67039.1      SLPYGANKEGIVWVATEGALNTPKDHIGTRNPNNNAAIVLQLPQGTTLPKGFYAEGSRGG
Q0Q468.1        SLPYGANKEGIVWVATEGALNTPKDHIGTRNPNNNAAIVLQLPQGTTLPKGFYAEGSRGG
ARI44802.1      SLPYGANKEGIVWVATEGALNTPKDHIGTRNPNNNAATVLQLPQGTTLPKGFYAEGSRGG
ARI44807.1      SLPYGANKEGIVWVATEGALNTPKDHIGTRNPNNNAATVLQLPQGTTLPKGFYAEGSRGG
ADE34820.1      SLPYGANKEGIVWVATEGALNTPKDHIGTRNPNNNAAVVLQLPQGTTLPKGFYAEGSRGG
Q3LZX4.1        SLPYGANKEGIVWVATEGALNTPKDHIGTRNPNNNAAIVLQLPQGTTLPKGFYAEGSRGG
AAZ41337.1      SLPYGANKEGIVWVATEGALNTPKDHIGTRNPNNNAAIVLQLPQGTTLPKGFYAEGSRGG
ADE34730.1      SLPYGANKEGIVWVATEGALNTPKDHIGTRNPNNNAAIVLQLPQGTTLPKGFYAEGSRGG
ADE34787.1      SLPYGANKEGIVWVATEGALNTPKDHIGTRNPNNNAAIVLQLPQGTTLPKGFYAEGSRGG
ATO98129.1      SLPYGANKEGIVWVATEGALNTPKDHIGTRNPNNNAATVLQLPQGTTLPKGFYAEGSRGG
ALK02467.1      SLPYGANKEGIVWVATEGALNTPKDHIGTRNPNNNAATVLQLPQGTTLPKGFYAEGSRGG
AGZ48815.1      SLPYGANKEGIVWVATEGALNTPKDHIGTRNPNNNAATVLQLPQGTTLPKGFYAEGSRGG
ATO98228.1      SLPYGANKEGIVWVATEGALNTPKDHIGTRNPNNNAATVLQLPQGTTLPKGFYAEGSRGG
ATO98190.1      SLPYGANKEGIVWVATEGALNTPKDHIGTRNPNNNAATVLQLPQGTTLPKGFYAEGSRGG
ATO98154.1      SLLYGANKEGIVWVATEGALNTPKDHIGTRNPNNNAATVLQLPQGTTLPKGFYAEGSRGG
AAR12990.1      SLPYGANKEGIVWVATEGALNTPKDHIGTRNPNNNAATVLQLPQGTTLPKGFYAEGSRGG
AAU04673.1      SLPYGANKEGIVWVATEGALNTPKDHIGTRNPNNNAATVLQLPQGTTLPKGFYAEGSRGG
ATO98142.1      SLPYGANKEGIVWVATEGALNTPKDHIGTRNPNNNAATVLQLPQGTTLPKGFYAEGSRGG
AAP82974.1      SLPYGANKEGIVWVATEGALNTPKDHIGTRNPNNNAATVLQLPQGTTLPKGFYAEGSRGG
ATO98117.1      SLPYGANKEGIVWVATEGALNTPKDHIGTRNPNNNVATVLQLPQGTTLPKGFYAEGSRGG
AAX16200.1      SLPYGANKEGIVWVATEGALNTPKDHIGTRNPNNNAATVLQLPQGTTLPKGFYAEGSRGG
AAV91640.1      SLPYGANKEGIVWVATEGALNTPKDHIGTRNPNNNAATVLQLPQGTTLPKGFYAEGSRGG
AAU04642.1      SLPYGANKEGIVWVATEGALNTPKDHIGTRNPNNNAATVLQLPQGTTLPKGFYAEGSRGG
AAV49739.1      SLPYGANKEGIVWVATEGALNTPKDHIGTRNPNNNGATVLQLPQGTTLPKGFYAEGSRGG
AAS48456.1      SLPYGANKEGIVWVATEGALNTPKDHIGTRNPNNNAATVLQLPQGTTLPKGFYAEGSRGG
AAU04658.1      SLPYGANKEGIVWVATEGALNTPKDHIGTRNPNNNAATVLQLPQGTTLPKGFYAEGSRGG
ACB69868.1      SLPYGANKEGIVWVATEGALNTPKDHIGTRNPNNNAATVLQLPQGTTLPKGFYAEGSRGG
QDF43833.1      SLPYGANKEGIVWVATEGALNTPKDHIGTRNPNNNAATVLQLPQGTTLPKGFYAEGSRGG
ACZ72205.1      SLPYGANKEGIVWVATEGALNTPKDHIGTRNPNNNAATVLQLPQGTTLPKGFYAEGSRGG
AAP33707.1      SLPYGANKEGIVWVATEGALNTPKDHIGTRNPNNNAATVLQLPQGTTLPKGFYAEGSRGG
ACZ72117.1      SLPYGANKEGIVWVATEGALNTPKDHIGTRNPNNNAATVLQLPQGTTLPKGFYAEGSRGG
AAT76155.1      SLPYGANKEGIVWVATEGALNTPKDHIGTRNPNNNAATVLQLPQGTTLPKGFYAEGSRGG
AGZ48841.1      SLPYGANKEGIVWVATEGALNTPKDHIGTRNPNNNAATVLQLPQGTTLPKGFYAEGSRGG
ATO98202.1      SLPYGANKEGIVWVATEGALNTPKDHIGTRNPNNNAATVLQLPQGTTLPKGFYAEGSRGG
ATO98166.1      SLPYGANKEGIIWVATEGALNTPKDHIGTRNPNNNAATVLQLPQGTTLPKGFYAEGSRGG
ACZ71986.1      SLPYGANKEGIVWVATEGALNTPKDHIGTRNPNNNAATVLQLPQGTTLPKGFYAEGSRGG
AAP30714.1      SLPYGANKEGIVWVATEGALNTPKDHIGTRNPNNNAATVLQLPQGTTLPKGFYAEGSRGG
AAR87518.1      SLPYGANKEGIIWVATEGALNTPKDHIGTRNPNNNAATVLQLPQGTTLPKGFYAEGSRGG
QDF43828.1      SLPYGANKEGIVWVATEGALNTPKDHIGTRNPNNNAAIVLQLPQGTTLPKGFYAEGSRGG
ATO98178.1      SLPYGANKEGIVWVATEGALNTPKDHIGTRNPNNNAATVLQLPQGTTLPKGFYAEGSRGG
NP_828858.1     SLPYGANKEGIVWVATEGALNTPKDHIGTRNPNNNAATVLQLPQGTTLPKGFYAEGSRGG
ACU31047.1      SLPYGANKEGIVWVATEGALNTPKDHIGTRNPNNNAAIVLQLPQGTTLPKGFYAEGSRGG
AID16719.1      SLPYGANKEGIVWVATEGALNTPKDHIGTRNPNNNAAIVLQLPQGTTLPKGFYAEGSRGG
QDF43818.1      SLPYGANKEGIVWVATEGALNTPKDHIGTRNPNNNAAIVLQLPQGTTLPKGFYAEGSRGG
AAZ67043.1      SLPYGANKEGIVWVATEGALNTPKDHIGTRNPNNNAAIVLQLPQGTTLPKGFYAEGSRGG
Q3I5I7.1        SLPYGANKEGIVWVATEGALNTPKDHIGTRNPNNNAAIVLQLPQGTTLPKGFYAEGSRGG
ADE34763.1      SLPYGANKEGIVWVATEGALNTPKDHIGTRNPNNNAAIVLQLPQGTTLPKGFYAEGSRGG
AHX37566.1      SLPYGANKEGIVWVATEGALNTPKDHIGTRNPNNNAAIVLQLPQGTTLPKGFYAEGSRGG
ARO76389.1      SLPYGANKEGIVWVATEGALNTPKDHIGTRNPNNNAAIVLQLPQGTTLPKGFYAEGSRGG
AGC74175.1      SLPYGANKEGIVWVAIEGALNTPKDHIGTRNPNNNAAIVLQLPQGTTLPKGFYAEGSRGG
AIA62338.1      SLPYGANKEGIVWVATEGALNTPKDHIGTRNPNNNAAIVLQLPQGTTLPKGFYAEGSRGG
AIA62328.1      SLPYGANKEGIVWVAIEGALNTPKDHIGTRNPNNNAAIVLQLPQGTTLPKGFYAEGSRGG
ACU31039.1      SLPYGANKQGIVWVATEGALNTPKDHIGTRNPNNNAAIVLQLPQGTTLPKGFYAEGSRGG
QDF43823.1      SLPYGANKEGIVWVATEGALNTPKDHIGTRNPNNNAAIVLQLPQGTTLPKGFYAEGSRGG
QIA48648.1      GLPYGANKEGIIWVATEGALNTPKDHIGTRNPNNNAAIVLQLPQGTALPKGFYAEGSRGG
QIQ54056.1      GLPYGANKEGIIWVATEGALNTPKDHIGTRNPNNNAAIVLQLPQGTALPKGFYAEGSRGG
QIA48621.1      GLPYGANKEGIIWVATEGALNTPKDHIGTRNPNNNAAIVLQLPQGTALPKGFYAEGSRGG
QIA48630.1      GLPYGANKEGIIWVATEGALNTPKDHIGTRNPNNNAAIVLQLPQGTALPKGFYAEGSRGG
QIA48639.1      GLPYGANKEGIIWVATEGALNTPKDHIGTRNPNNNAAIVLQLPQGTALPKGFYAEGSRGG
AVP78038.1      GLPYGANKEGIIWVATEGALNTPKDHIGTRNPANNAAIVLQLPQGTTLPKGFYAEGSRGG
AVP78049.1      GLPYGANKEGIIWVATEGALNTPKDHIGTRNPANNAAIVLQLPQGTTLPKGFYAEGSRGG
QIS61476.1      GLPYGANKDGIIWVATEGALNTPKDHIGTRNPANNAAIVLQLPQGTTLPKGFYAEGSRGG
QIQ50109.1      GLPYGANKDGIIWVATEGALNTPKDHIGTRNPANNAAIVLQLPQGTTLPKGFYAEGSRGG
QHR63308.1      GLPYGANKDGIIWVATEGALNTPKDHIGTRNPANNAAIVLQLPQGTTLPKGFYAEGSRGG
QIS30122.1      GLPYGANKDGIIWVATEGALNTPKDHIGTRNPANNAAIVLQLPQGTTLPKGFYAEGSRGG
QIJ96530.1      GLPYGANKDGIIWVATEGALNTPKDHIGTRNPANNAAIVLQLPQGTTLPKGFYAEGSRGG
QIQ08827.1      GLPYGANKDGIIWVATEGALNTPKDHIGTRNPANNAAIVLQLPQGTTLPKGFYAEGSRGG
QIS30182.1      GLPYGANKDGIIWVATEGALNTPKDHIGTRNPANNAAIVLQLPQGTTLPKGFYAEGSRGG
QIH45050.1      GLPYGANKDGIIWVATEGALNTPKDHIGTRNPANNAAIVLQLPQGTTLPKGFYAEGSRGG
QIS30511.1      GLPYGANKDGIIWVATEGALNTPKDHIGTRNPANNAAIVLQLPQGTTLPKGFYAEGSRGG
QHW06046.1      GLPYGANKDGIIWVATEGALNTPKDHIGTRNPANNAAIVLQLPQGTTLPKGFYAEGSRGG
QIM47464.1      GLPYGANKDGIIWVATEGALNTPKDHIGTRNPANNAAIVLQLPQGTTLPKGFYAEGSRGG
QIS30192.1      GLPYGANKDGIIWVATEGALNTPKDHIGTRNPANNAAIVLQLPQGTTLPKGFYAEGSRGG
QII57305.1      GLPYGANKDGIIWVATEGALNTPKDHIGTRNPANNAAIVLQLPQGTTLPKGFYAEGSRGG
QIE07458.1      GLPYGANKDGIIWVATEGALNTPKDHIGTRNPANNAAIVLQLPQGTTLPKGFYAEGSRGG
QIS61010.1      GLPYGANKDGIIWVATEGALNTPKDHIGTRNPANNAAIVLQLPQGTTLPKGFYAEGSRGG
QIS30062.1      GLPYGANKDGIIWVATEGALNTPKDHIGTRNPANNAAIVLQLPQGTTLPKGFYAEGSRGG
QIQ50129.1      GLPYGANKDGIIWVATEGALNTPKDHIGTRNPANNAAIVLQLPQGTTLPKGFYAEGSRGG
QIS30552.1      GLPYGANKDGIIWVATEGALNTPKDHIGTRNPANNAAIVLQLPQGTTLPKGFYAEGSRGG
QIS60866.1      GLPYGANKDGIIWVATEGALNTPKDHIGTRNPANNAAIVLQLPQGTTLPKGFYAEGSRGG
QHO62884.1      GLPYGANKDGIIWVATEGALNTPKDHIGTRNPANNAAIVLQLPQGTTLPKGFYAEGSRGG
QIS30502.1      GLPYGANKDGIIWVATEGALNTPKDHIGTRNPANNAAIVLQLPQGTTLPKGFYAEGSRGG
QIC50514.1      GLPYGANKDGIIWVATEGALNTPKDHIGTRNPANNAAIVLQLPQGTTLPKGFYAEGSRGG
BCB97908.1      GLPYGANKDGIIWVATEGALNTPKDHIGTRNPANNAAIVLQLPQGTTLPKGFYAEGSRGG
QIS60878.1      GLPYGANKDGIIWVATEGALNTPKDHIGTRNPANNAAIVLQLPQGTTLPKGFYAEGSRGG
YP_009724397.2  GLPYGANKDGIIWVATEGALNTPKDHIGTRNPANNAAIVLQLPQGTTLPKGFYAEGSRGG
QIQ49839.1      GLPYGANKDGIIWVATEGALNTPKDHIGTRNPANNAAIVLQLPQGTTLPKGFYAEGSRGG
YP_003858591.1  ALPYGANKDGIVWVATEGALNTPKDHIGTRNPNNNAAIVIQLPQGTTLPKGFYAEGSRGG
APO40586.1      ALPYGANKEGIVWVATEGALNTPKEHIGTRNPNNNAAIVLQLPQGTSLPKGFYAEGSRGG
```

```
                             190       200       210       220       230       240
AAS01074.1          SQASSRSSSRXXXNSRNSTPGSSRGNSPARMASGGGETALALLLLDRLNQLESKVSGKGQ
ABD75315.1          SQASSRSSSRSRGNSRTSTPGSSRGNSPARVASGGGETALALLLLDRLNQLESKVSGKGQ
AIA62298.1          SQASSRSSSRSRGNSRTSTPGSSRGNSPARVASGGGETALALLLLDRLNQLESKVSGKGQ
ABG47067.1          SQASSRSSSRSRGNSRTSTPGSSRGNSPARVASGGGETALALLLLDRLNQLESKVSGKGQ
ANA96035.1          SQASSRSSSRSRGNSRTSTPGSSRGNSPARVASGGGETALALLLLDRLNQLESKVSGKGQ
AIA62285.1          SQASSRSSSRSRGNSRISTPGSSRGNSPARVASGGGETALALLLLDRLNQLESKVSGKGQ
ASO66816.1          SQASSRSSSRSRGNSRTSTPGSSRGNSPARVASGGGETALALLLLDRLNQLESKVSGKGQ
AKZ19095.1          SQASSRSSSRSRGNSRNSTPGSSRGTSPARIASGGGETALALLLLDRLNQLESKVSGKGQ
AKZ19084.1          SQASSRSSSRSRGNSRNSTPGSSRGTSPARIASGGGETALALLLLDRLNQLESKVSGKGQ
AIA62318.1          SQASSRSSSRSRGNSRNSTPGSSRGNSPARMASGGGETALALLLLDRLNQLESKVSGKGQ
AGC74169.1          SQASSRSSSRSRGNSRNSTPGSSRGNSPARMASGGGETALALLLLDRLNQLESKVSGKGQ
AAZ67039.1          SQASSRSSSRSRGNSRNSTPGSSRGNSPARMASGSGETALALLLLDRLNQLESKVSGKGQ
Q0Q468.1            SQASSRSSSRSRGNSRNSTPGSSRGNSPARMASGSGETALALLLLDRLNQLESKVSGKGQ
ARI44802.1          SQASSRSSSRSRGNSRNSTPGSSRGNSPARMASGGGETALALLLLDRLNQLESKVSGKGQ
ARI44807.1          SQASSRSSSRSRGNSRNSTPGSSRGNSPARMAIGGGETALALLLLDRLNQLESKVSGKGQ
ADE34820.1          SQSSSRSSSRSRGNSTNSTPGSSRGSSPARLASGGGETALALLLLDRLNQLESKVSGKGQ
Q3LZX4.1            SQSSSRSSSRSRGNSRNSTPGSSRGSSPARLASGGGETALALLLLDRLNQLESKVSGKGQ
AAZ41337.1          SQSSSRSSSRSRGNSRNSTPGSSRGSSPARLASGGGETALALLLLDRLNQLESKVSGKGQ
ADE34730.1          SQSSSRSSSRSRGNSTNSTPGSSRGSSPARLASGGGETALALLLLDRLNQLESKVSGKGQ
ADE34787.1          SQSSSRSSSRSRGNSTNSTPGSSRGSSPARLASGGGETALALLLLDRLNQLESKVSGKGQ
ATO98129.1          SQASSRSSSRSRGDSRNSTPGSSRGNSPARMASGGGETALALLLLDRLNQLESKVSGKGQ
ALK02467.1          SQASSRSSSRSRGNSRNSTPGSSRGNSPARMASGGGETALALLLLDRLNQLESKVSGKGQ
AGZ48815.1          SQASSRSSSRSRGNSRNSTPGSSRGNSPARMASGGGETALALLLLDRLNQLESKVSGKGQ
ATO98228.1          SQASSRSSSRSRGNSRNSTPGSSRGNSPARMASGGGETALALLLLDRLNQLESKVSGKGQ
ATO98190.1          SQASSRSSSRSRGNSRNSTPGSSRGNSPARMASGGGETALALLLLDRLNQLESKVSGKGQ
ATO98154.1          SQASSRSSSRSRGNSRNSTPGSSRGNSPARMASGGGETALALLLLDRLNQLESKVSGKGQ
AAR12990.1          SQASSRSSSRSRGNSRNSTPGSSRGNSPARMASGGGETALALLLLDRLNRLESKVSGKGQ
AAU04673.1          SQASSRSSSRSRGNSRNSTPGSSRGNSPARMASGGGETALALLLLDRLNQLESKVSGKGQ
ATO98142.1          SQASSRSSSRSRGNSRNSTPGSSRGNSPARMASGGGETALALLLLDRLNQLESKVSGKGQ
AAP82974.1          SQASSRSSSRSRGNSRNSTPGSSRGNSPARMASGGGETALALLLLDRLNQLESKVSGKGQ
ATO98117.1          SQASSRSSSRSRGNSRNSTPGSSRGNSPARMASGGGETALALLLLDRLNQLESKVSGKGQ
AAX16200.1          SQASSRSSSRSRGNSGNSTPGSSRGNSPARMASGGGKTALALLLLDRLNQLESKVSGKGQ
AAV91640.1          SQASSRSSSRSRGNSRNSTPGSSRGNSPARMASGGGETALALLLLDRLNQLESKVSGKGQ
AAU04642.1          SQASSRSSSRSRGNSRNSTPGSSRGNSPARMASGGGETALALLLLDRLNQLESKVSGKGQ
AAV49739.1          SQASSRSSSRSRGNSRNSTPGSSRGNSPARMASGGGETALALLLLDRLNQLESKVSGKGQ
AAS48456.1          SQASSRSSSRSRGNSRNSTPGSSRGNSPARMASGGGETALALLLLDRLNQLESKVSGKGQ
AAU04658.1          SQASSRSSSRSRGNSRNSTPGSSRGNSPARMASGGGETALALLLLDRLNQLESKVSGKGQ
ACB69868.1          SQASSRSSSRSRGNSRNSTPGSSRGNSPARMASGGGETALALLLLDRLNQLESKVSGKGQ
QDF43833.1          SQASSRSSSRSRGNSRNSTPGSSRGNSPARMASGGGETALALLLLDRLNQLESKVSGKGQ
ACZ72205.1          SQASSRSSSRSRGNSRNSTPGSSRGNSPARMASGGGETALALLLLDRLNQLESKVSGKGQ
AAP33707.1          SQASSRSSSRSRGNSRNSTPGSSRGNSPARMASGGGETALALLLLDRLNQLESKVSGKGQ
ACZ72117.1          SQASSRSSSRSRGNSRNSTPGSSRGNSPARMASGGGETALALLLLDRLNQLESKVSGKGQ
AAT76155.1          SQASSRSSSRSRGNSRNSTPGSSRGNSPARMASGGGETALALLLLDRLNQLESKVSGKGQ
AGZ48841.1          SQASSRSSSRSRGNSRNSTPGSSRGNSPARMASGGGETALALLLLDRLNQLESKVSGKGQ
ATO98202.1          SQASSRSSSRSRGNSRNSTPGSSRGNSPARMASGGGETALALLLLDRLNQLESKVSGKGQ
ATO98166.1          SQASSRSSSRSRGNSRNSTPGSSRGNSPARMASGGGETALALLLLDRLNQLESKVSGKGQ
ACZ71986.1          SQASSRSSSRSRGNSRNSTPGSSRGISPARMASGGGETALALLLLDRLNQLESKVSGKGQ
AAP30714.1          SQASSRSSSRSRCNSRNSTPGSSRGNSPARMASGGGETALALLLLDRLNQLESKVSGKGQ
AAR87518.1          SQASSRSSSRSRGNSRNSTPGSSRGNSPARMASGGGETALALLLLDRLNQLESKVSGKGQ
QDF43828.1          SQASSRSSSRSRGNSRNSTPGSSRGNSPARMASGGGETALALLLLDRLNQLESKVSGKGQ
ATO98178.1          SQASSRSSSRSRGNSRNSTPGSSRGNSPARMASGGGETALALLLLDRLNQLESKVSGKGQ
NP_828858.1         SQASSRSSSRSRGNSRNSTPGSSRGNSPARMASGGGETALALLLLDRLNQLESKVSGKGQ
ACU31047.1          SQASSRSSSRSRGNSRNSTPGSSRGNSPARLASGGGETALALLLLXRLNQLESKVSGKGQ
AID16719.1          SQASSRSSSRSRGNSRNSTPGSSRGNSPARLASGGGETALALLLLDRLNQLESKVSGKGQ
QDF43818.1          SQASSRSSSRSRGNSRNSTPGSSRGNSPARMASGGGETALALLLLDRLNQLESKVSGKGQ
AAZ67043.1          SQASSRSSSRSRGNSRNSTPGSSRGNSPARMASGGGETALALLLLDRLNQLESKVSGRSQ
Q3I5I7.1            SQASSRSSSRSRGNSRNSTPGSSRGNSPARMASGGGETALALLLLDRLNQLESKVSGRSQ
ADE34763.1          SQASSRSSSRSRGNSRNSTPGSSRGNSPARLASGGGETALALLLLDRLNQLESKVSGKGQ
AHX37566.1          SQASSRSSSRSRGNSRNSTPGSSRGNSPARMASGGGETALALLLLDRLNQLESKVSGKGQ
ARO76389.1          SQASSRSSSRSRGNSRNSTPGSSRGNSPARMASGGGETALALLLLDRLNQLESKVSGKGQ
AGC74175.1          SQASSRSSSRSRGNSRNSTPGSSRGNSPARMASGGGETALALLLLDRLNQLESKVSGKGQ
AIA62338.1          SQASSRSSSRSRGNSRNSTPGSSRGNSPARMASGGGETALALLLLDRLNQLESKVSGKGQ
AIA62328.1          SQASSRSSSRSRGNSRNSTPGSSRGNSPARLASGGGETALALLLLDRLNQLESKVSGKGQ
ACU31039.1          SQASSRSSSRSRGNSRNSTPGSSRGNSPARMASGGGETALALLLLDRLNQLESKVSGKGQ
QDF43823.1          SQASSRSSSRSRGNSRNSTPGSSRGNSPARTASGGGETALALLLLDRLNQLESKVSGKGQ
QIA48648.1          SQASSRSSSRSRNSSRNSTPGSSRGTSPARIAGNGGDAALALLLLDRLNALESKMSGKGS
QIQ54056.1          SQASSRSSSRSRNSSRNSTPGSSRGTSPARIAGNGGDAALALLLLDRLNALESKMSGKGS
QIA48621.1          SQASSRSSSRSRNSSRNSTPGSSRGTSPARIAGNGGDAALALLLLDRLNALESKMSGKGS
QIA48630.1          SQASSRSSSRSRNSSRNSTPGSSRGTSPARIAGNGGDAALALLLLDRLNALESKMSGKGS
QIA48639.1          SQASSRSSSRSRNSSRNSTPGSSRGTSPARIAGNGGDAALALLLLDRLNALESKMSGKGS
AVP78038.1          SQASSRSSSRSRNSSRNSTPGSSRGTSPARMAGNGGDTALALLLLDRLNQLENKVSGKGQ
AVP78049.1          SQASSRSSSRSRNSSRNSTPGSSRGTSPARMAGNGGDTALALLLLDRLNQLENKISGKGQ
QIS61476.1          SQASSRSSSRSRNSSRNSTPGSSRGTSPARMAGNGGDAALALLLLDRLNQLESKMSGKGQ
QIQ50109.1          SQASSRSSSRSRNSSRNSTPGSSRGTSPARMAGNGGDAALALLLLDRLNQLESKMSGKGQ
QHR63308.1          SQASSRSSSRSRNSSRNSTPGSSRGTSPARMAGNGSDAALALLLLDRLNQLESKMSGKGQ
QIS30122.1          SQASSRSSSRSRNSSRNSTPGSSRGTSPARMAGNGGDAALALLLLDRLNQLESKMSGKGQ
QIJ96530.1          SQASSRSSSRSRNSSRNSTPGSSRGTSPARMAGNGSDAALALLLLDRLNQLESKMSGKGQ
QIQ08827.1          SQASSRSSSRSRNSSRNSTPGSSKRTSPARMAGNGGDAALALLLLDRLNQLESKMSGKGQ
QIS30182.1          SQASSRSSSRSRNSSRNSTPGSSRGTSPARMAGNGGDAALALLLLDRLNQLESKMSGKGQ
QIH45050.1          SQASSRSSSRSRNSSRNSTPGSSRGTSPARMAGNGGDAALALLLLDRLNQLESKMSGKGQ
QIS30511.1          SQASSRSSSRSRNSSRNSTPGSSRGTSPARMAGNGGDAALALLLLDRLNQLESKMSGKGQ
QHW06046.1          SQASSRSSSRSRNSLRNSTPGSSRGTSPARMAGNGGDAALALLLLDRLNQLESKMSGKGQ
QIM47464.1          SQASSRSSSRSRNSSRNLTPGSSRGTSPARMAGNGGDAALALLLLDRLNQLESKMSGKGQ
QIS30192.1          SQASSRSSSRSRNSSRNSTPGSSRGTSPARMAGNGGDAALALLLLDRLNQLESKMSGKGQ
QII57305.1          SQASSRSSSRSRNSSRNSTPGSSRGTSPARMAGNGGDAALALLLLDRLNQLESKMSGKGQ
QIE07458.1          SQASSRSSSRSRNSSRNSTPGSSRGTSPARMAGNGGDAALALLLLDRLNQLESKMSGKGQ
QIS61010.1          SQASSRSSSRSRNSSRNSTPGSSRGTSPARMAGNGGDAALALLLLDRLNQLESKMSGKGQ
QIS30062.1          SQASSRSSSRSRNSSRNSTPGSSRGTSPARMAGNGGDAALALLLLDRLNQLESKMSGKCQ
QIQ50129.1          SQASSCSSSRSRNSSRNSTPGSSRGTSPARMAGNGGDAALALLLLDRLNQLESKMSGKGQ
QIS30552.1          SQASSRSSSRSRNSSRNSTPGSSRGTSPARMAGNGGDAALALLLLDRLNQLESKMSGKGQ
QIS60866.1          SQASSRSSSRSRNSSRNSTPGSSRGTSPGRMAGNGGDAALALLLLDRLNQLESKMSGKGQ
QHO62884.1          SQASSRSSSRSRNSXRNSTPGSSRGTSPARMAGNGGDAALALLLLDRLNQLESKMSGKGQ
QIS30502.1          SQASSRSSSRSRNSSRNSTPGSSRGTSPARMAGNGGDAALALLLLDRLNQLESKMSGKGQ
QIC50514.1          SQASSRSSSRSRNSSRNSTPGSNRGTSPARMAGNGGDAALALLLLDRLNQLESKMSGKGQ
BCB97908.1          SQASSRSSSRSRNSSRNSTPGSSRGTSPARMAGNGGDAALALLLLDRLNQLESKMSGKGQ
QIS60878.1          SQASSRSSSRSRNSSRNSTPGSSRGTSPARMAGNGGDAALALLLLDRLNQLETKMSGKGQ
YP_009724397.2      SQASSRSSSRSRNSSRNSTPGSSRGTSPARMAGNGGDAALALLLLDRLNQLESKMSGKGQ
QIQ49839.1          SQASSRSSSRSRNSSRNSTPGSSKGTSPARMAGNGGDAALALLLLDRLNQLESKMSGKGQ
YP_003858591.1      SQASSRSNSRSRGNSRNSTPSSSRGSSPAR.MAAGGDTALALLLLDRLNQLESKVSGKTP
APO40586.1          SQTASRSSSRSRGNSRNSTPSSSRGSSPARNLQAGGDTALALLLLDRLNQLESKVSGKTQ
```

```
                          250       260
                            .         .
AAS01074.1      QQQGXTVTKKSAAEASKKPRQK......................................
ABD75315.1      QQQGQTVTKKSTSEASKKPRQKRTATKQYNVTQAFGRRGPDQTQGNFGDQELIRQGTDYK
AIA62298.1      QQQGQTVTKKSASEASKKPRQKRTATKQYNVTQAFGRRGPDQTQGNFGDQELIRQGTDYK
ABG47067.1      QQQGQTVTKKSASEASKKPRQKRTATKQYNVTQAFGRRGPDQTQGNFGDQELIRQGTDYK
ANA96035.1      QQQGQTVTKKSASEASKKPRQKRTATKQYNVTQAFGRRGPDQTQGNFGDQELIRQGTDYK
AIA62285.1      QQQGQTVTKKSASEASKKPRQKRTATKQYNVTQAFGRRGPDQTQGNFGDQELIRQGTDYK
ASO66816.1      QQQGQTVTKKSASEASKKPRQKRTATKQYNVTQAFGRRGPDQTQGNFGDQELIRQGTDYK
AKZ19095.1      QQQGQTVTKKSAAEASKKPRQKRTATKQYNVTQAFGRRGPDQTQGNFGDQDLIRQGTDYK
AKZ19084.1      QQQGQTVTKKSAAEASKKPRQKRTATKQYNVTQAFGRRGPDQTQGNFGDQDLIRQGTDYK
AIA62318.1      QQQGQTVTKKSAAEASKKPRQKRTATKSYNVTQAFGRRGPEQTQGNFGDQELIRQGTDYK
AGC74169.1      QQQGQTVTKKSAAEASKKPRQKRTATKSYNVTQAFGRRGPEQTQGNFGDQDLIRQGTDYK
AAZ67039.1      QQQGQTVTKKSAAEASKKPRQKRTATKSYNVTQAFGRRGPEQTQGNFGDQDLIRQGTDYK
Q0Q468.1        QQQGQTVTKKSAAEASKKPRQKRTATKSYNVTQAFGRRGPEQTQGNFGDQDLIRQGTDYK
ARI44802.1      Q.QGQTVTKKSAAEASKKPRQKRTATKQYNVTQAFGRRGPEQTQGNFGDQELIRQGTDYK
ARI44807.1      Q.QGQTVTKKSAAEASKKPRQKRTATKQYNVTQAFGRRGPEQTQGNFGDQELIRQGTDYK
ADE34820.1      QQPGQTVTKKSAAEASKKPRQKRTATKQYNVTQAFGRRGPEQTQGNFGDQELIRQGTDYK
Q3LZX4.1        QQPGQTVTKKSAAEASKKPRQKRTATKQYNVTQAFGRRGPEQTQGNFGDQELIRQGIDYK
AAZ41337.1      QQPGQTVTKKSAAEASKKPRQKRTATKQYNVTQAFGRRGPEQTQGNFGDQELIRQGTDYK
ADE34730.1      QQPGQTVTKKSAAEASKKPRQKRTATKQYNVTQAFGRRGPEQTQGNFGDQELIRQGTDYK
ADE34787.1      QQPGQTVTKKSAAEASKKPRQKRTATKQYNVTQAFGRRGPEQTQGNFGDQQLIRQGTDYK
ATO98129.1      QQQGQTVTKKSAAEASKKPRQKRTATKQYNVTQAFGRRGPEQTQGNFGDQDLIRQGTDYK
ALK02467.1      QQQGQTVTKKSAAEASKKPRQKRTATKQYNVTQAFGRRGPEQTQGNFGDQDLIRQGTDYK
AGZ48815.1      QQQGQTVTKKSAAEASKKPRQKRTATKQYNVTQAFGRRGPEQTQGNFGDQDLIRQGTDYK
ATO98228.1      QQQGQTVTKKSAAEASKKPRQKRTATKQYNVTQAFGRRGPEQTQGNFGDQDLIRQGTDYK
ATO98190.1      QQQGQTVTKKSAAEASKKPRQKRTATKQYNVTQAFGRRGPEQTQGNFGDQDLIRQGTDYK
ATO98154.1      QQQGQTVTKKSAAEASKKPRQKRTATKQYNVTQAFGRRGPEQTQGNFGDQDLIRQGTDYK
AAR12990.1      QQQGQTVTKKSAAEASKKPRQKRTATKQYNVTQAFGRRGPEQTQGNFGDQDLIRQGTDYK
AAU04673.1      QQQGQTVTKKSAAEASKKPRQKRTATKQYNVTQAFGRRGPEQTQGNFGDQDLIRQGIDYK
ATO98142.1      QQQGQTVTKKSAAEASKKPRQKRTATKQYSVTQAFGRRGPEQTQGNFGDQDLIRQGTDYK
AAP82974.1      QQQGQTVTKKSAAEASKKPRQKRTATKQYNVTQAFGRRGPEQTQGNFGDQDLIRQGTDYK
ATO98117.1      QQQGQTVTKKSAAEASKKPRQKRTATKQYNVTQAFGRRGPEQTQGNFGDQDLIRQGTDYK
AAX16200.1      QQQGQTVTKKSAAEASKKPRQKRTATKQYNVTQAFGRRGPEQTQGNFGDQDLIRQGTDYK
AAV91640.1      QQQGQTVTKKSAAEASKKPRQKRTATKQYNVTQAFGRRGPEQTQGNFGDQDLIRQGTDYK
AAU04642.1      QQQGQTVTKKSAAEASKKPRQKRTATKQYNVTQAFGRRGPEQTQGNFGDQDLIRQGTDYK
AAV49739.1      QQQGQTVTKKSAAEASKKPRQKRTATKQYNVTQAFGRRGPEQTQGNFGDQDLIRQGTDYK
AAS48456.1      QQQGQTVTKKSAAEASKKPRQKRTATKQYNVTQAFGRRGPEQTQGNFGDQDLIRQGTDYK
AAU04658.1      QQQGQTVTKKSAAEASKKPRQKRTATKQYNVTQAFGRRGPEQTQGNFGDQDLIRQGTDYK
ACB69868.1      QQQGQTVTKKSAAEASKKPRQKRTATKQYNVTQAFGRRGPEQTQGNFGDQDLIRQGTDYK
QDF43833.1      QQQGQTVTKKSAAEASKKPRQKRTATKQYNVTQAFGRRGPEQTQGNFGDQDLIRQGTDYK
ACZ72205.1      QQQGQTVTKKSAAEASKKPRQKRTATKQYNVTQAFGRRGPEQTQGNFGDQDLIRQGXDYK
AAP33707.1      QQQGQTVTKKSAAEASKKPRQKRTATKQYNVTQAFGRRGPEQTQGNFGDQDLIRQGTDYK
ACZ72117.1      QQQGQTVTKKSAAEASKKPRQKRTATKQYNVTQAFGRRGPEQTQGNFGDQDLIRQGTDYK
AAT76155.1      QQQGQTVTKKSAAEASKKPRQKRTATKQYNVTQAFGRRGPEQTQGNFGDQDLIRQGTDYK
AGZ48841.1      QQQGQTVTKKSAAEASRKPRQKRTATKQYNVTQAFGRRGPEQTQGNFGDQDLIRQGTDYK
ATO98202.1      QQQGQTVTKKSAAEASKKPRQKRTATKQYNVTQAFGRRGPEQTQGNFGDQDLIRQGTDYK
ATO98166.1      QQQGQTVTKKSAAEASKKPRQKRTATKQYNVTQAFGRRGPEQTQGNFGDQDLIRQGTDYK
ACZ71986.1      QQQGQTVTKKSAAEASKKPRQKRTATKQYNVTQAFGRRGPEQTQGNFGDQDLIRQGTDYK
AAP30714.1      QQQGQTVTKKSAAEASKKPRQKRTATKQYNVTQAFGRRGPEQTQGNFGDQDLIRQGTDYK
AAR87518.1      QQQGQTVTKKSAAEASKKPRQKRTATKQYNVTQAFGRRGPEQTQGNFGDQDLIRQGTDYK
QDF43828.1      QQQGQTVTKKSAAEASKKPRQKRTATKQYNVTQAFGRRGPEQTQGNFGDQDLIRQGTDYK
ATO98178.1      QQQGQTVTKKSAAEASKKPRQKRTATKAYNVTQAFGRRGPEQTQGNFGDQDLIRQGTDYK
NP_828858.1     QQQGQTVTKKSAAEASKKPRQKRTATKQYNVTQAFGRRGPEQTQGNFGDQDLIRQGTDYK
ACU31047.1      QQPGQTVTKKSAAEASKKPRQKRTATKQYNVTQAFGRRGPEQTQGNFGDQDLIRQGTDYK
AID16719.1      QQPGQTVTKKSAAEASKKPRQKRTATKQYNVTQAFGRRGPEQTQGNFGDQELIRQGTDYK
QDF43818.1      QQQGQTVTKKSAAEASKKPRQKRTATKQYNVTQAFGRRGPEQTQGNFGDQELIRQGTDYK
AAZ67043.1      QQQGQTVTKKSAAEASKKPRQKRTATKQYNVTQAFGRRGPEQTQGNFGDQELVRQGTDYK
Q3I5I7.1        QQQGQTVTKKSAAEASKKPRQKRTATKQYNVTQAFGRRGPEQTQGNFGDQELIRQGTDYK
ADE34763.1      QQQGQTVTKKSAAEASKKPRQKRTATKQYNVTQAFGRRGPEQTQGNFGDQDLIRQGTDYK
AHX37566.1      QQQGQTVTKKSAAEASKKPRQKRTATKAYNVTQAFGRRGPEQTQGNFGDQELIRQGTDYK
ARO76389.1      QQQGQTVTKKSAAEASKKPRQKRTATKQYNVTQAFGRRGPEQTQGNFGDQELIRQGTDYK
AGC74175.1      QQQGQTVTKKSAAEASKKPRQKRTATKQYNVTQAFGRRGPEQTQGNFGDQELIRQGTDYK
AIA62338.1      QQQGQTVTKKSAAEASKKPRQKRTATKVYNVTQAFGRRGPEQTQGNFGDQELIRQGTDYK
AIA62328.1      QQQGQTVTKKSAAEASKKPRQKRTATKQYNVTQAFGRRGPEQTQGNFGDQDLTRQGTDYK
ACU31039.1      QQQGQTVTKKSAAEASKKPRQKRTATKQYNVTQAFGRRGPEQTQGNFGDQDLIRQGTDYK
QDF43823.1      QQQGQTVTKKSAAEASKKPRQKRTATKQYNVTQAFGRRGPEQTQGNFGDQDLIRQGTDYK
QIA48648.1      QQQSQTVTKKSAAEASKKPRQKRTATKQYNVTQAFGRRGPEKTQGNFGDKELIRQGTEYK
QIQ54056.1      QQQSQTVTKKSAAEASKKPRQKRTATKQYNVTQAFGRRGPEQTQGNFGDQELIRQGTEYK
QIA48621.1      QQQSQTVTKKSAAEASKKPRQKRTATKQYNVTQAFGRRGPEQTQGNFGDQELIRQGTEYK
QIA48630.1      QQQSQTVTKKSAAEASKKPRQKRTATKQYNVTQAFGRRGPEQTQGNFGDQELIRQGTEYK
QIA48639.1      QKQSQTVTKKSAAEASKKPRQKRTATKQYNVTQAFGRRGPEQTQGNFGDQELIRQGTEYK
AVP78038.1      QQQGQTVTKKSAAEASKKPRQKRTATKQYNVTQAFGRRGPEQTQGNFGDQELIRQGTDYK
AVP78049.1      QQQGQTVTKKSAAEASKKPRQKRTATKQYNVTQAFGRRGPEQTQGNFGDQELIRQGTDYK
QIS61476.1      QQQGQTVTKKSAAEASKKPRQKRTATKAYNVTQAFGRRGPEQTQGNFGDQELIRQGTDXX
QIQ50109.1      QQQGQTVTKKSAAEASKKPRQKRTATKAYNVTQAFGRRGPEQTQGNFGDQELIRQGTDYK
QHR63308.1      QQQSQTVTKKSAAEASKKPRQKRTATKQYNVTQAFGRRGPEQTQGNFGDQELIRQGTDYK
QIS30122.1      QQQGQTVTKKSAAEASKKPRQKRTATKAYNVTQAFGRRGPEQTQGNFGDQELIRQGTDYK
QIJ96530.1      QQQGQTVTKKSAAEASKKPRQKRTATKAYNVTQAFGRRGPEQTQGNFGDQELIRQGTDYK
QIQ08827.1      QQQGQTVTKKSAAEASKKPRQKRTATKAYNVTQAFGRRGPEQTQGNFGDQELIRQGTDYK
QIS30182.1      QQQGQTVTKKSAAEASKKPRQKRTATKAYNVTQAFGRRGPEQTQGNFGDQELIRQGTDYK
QIH45050.1      QQQGQTVTKKSAAEASKKPRQKRTATKAYNVTQAFGRRGPEQTQGNFGDQELIRQGTDYK
QIS30511.1      QQQGQTVTKKSAAEASKKPRQKRTATKAYNVTQAFGRRGPEQTQGNFGDQELIRQGTDYK
QHW06046.1      QQQGQTVTKKSAAEASKKPRQKRTATKAYNVTQAFGRRGPEQTQGNFGDQELIRQGTDYK
QIM47464.1      QQQGQTVTKKSAAEASKKPRQKRTATKAYNVTQAFGRRGPEQTQGNFGDQELIRQGTDYK
QIS30192.1      QQQGQTVTKKSAAEASKKPRQKRTATKAYNVTQAFGRRGPEQTQGNFGDQELIRQGTDYK
QII57305.1      QQQGQTVTKKSAAEASKKPRQKRTATKAYNVTQAFGRRGPEQTQGNFGDQELIRQGTDYK
QIE07458.1      QQQGQTVTKKSAAEASKKPRQKRTATKAYNVTQAFGRRGPEQTQGNFGDQELIRQGTDYK
QIS61010.1      QQQGQTVTKKSAAEASKKPRQKRTATKAYNVIQAFGRRGPEQTQGNFGDQELIRQGTDYK
QIS30062.1      QQQGQTVTKKSAAEASKKPRQKRTATKAYNVTQAFGRRGPEQTQGNFGDQELIRQGTDYK
QIQ50129.1      QQQGQTVTKKSAAEASKKPRQKRTATKAYNVTQAFGRRGPEQTQGNFGDQELIRQGTDYK
QIS30552.1      QQQGQTVTKKSAAEASKKPRQKRTATKAYNVTQAFGRRGPEQTQGNFGDQELIRQGTDYK
QIS60866.1      QQQGQTVTKKSAAEASKKPRQKRTATKAYNVTQAFGRRGPEQTQGNFGDQELIRQGTDYK
QHO62884.1      QQQGQTVTKKSAAEASKKPRQKRTATKAYNVTQAFGRRGPEQTQGNFGDQELIRQGTDYK
QIS30502.1      QQQGQTVTKKSAAEASKKPRQKRTATKAYNVTQAFGRRGPEQTQGNFGDQELIRQGTDYK
QIC50514.1      QQQGQTVTKKSAAEASKKPRQKRTATKAYNVTQAFGRRGPEQTQGNFGDQELIRQGTDYK
BCB97908.1      QQQGQTVTKKSAAEASKKPRQKRTATKAYNVTQAFGRRGPEQTQGNFGDHELIRQGTDYK
QIS60878.1      QQQGQTVTKKSAAEASKKPRQKRTATKAYNVTQAFGRRGPEQTQGNFGDQELIRQGTDYK
YP_009724397.2  QQQGQTVTKKSAAEASKKPRQKRTATKAYNVTQAFGRRGPEQTQGNFGDQELIRQGTDYK
QIQ49839.1      QQQGQTVTKKSAAEASKKPRQKRTATKAYNVTQAFGRRGPEQTQGNFGDQELIRQGTDYK
YP_003858591.1  Q.QSQVVTKKTAAEASKKPRQKRTATKAYNVTQAFGRRGPEPTQGNFGDQELIRLGTDYK
APO40586.1      QQQPQVVTRKSASEASKKPRQKRTATKQYNVTQAFGRRGPEQTQGNFGDQELIRLGTDYK
```

```
AAS01074.1       ...............................................................
ABD75315.1       HWPQIAQFAPSASAFFGMSRISMEVTPSGTWLIYHGAIKLDDKDPQFKDNVILLNKHIDA
AIA62298.1       HWPQIAQFAPSASAFFGMSRIGMEVTPSGTWLTYHGAIKLDDKDPQFKDNVILLNKHIDA
ABG47067.1       HWPQIAQFAPSASAFFGMSRISMEVTPSGTWLTYHGAIKLDDKDPQFKDNVILLNKHIDA
ANA96035.1       HWPQIAQFAPSASAFFGMSRIGMEVTPSGTWLTYHGAIKLDDKDPQFKDNVILLNKHIDA
AIA62285.1       HWPQIAQFAPSASAFFGMSRIGMEVTPSGTWLTYHGAIKLDDKDPQFKDNVILLNKHIDA
ASO66816.1       HWPQIAQFAPSASAFFGMSRIGMEVTPSGTWLTYHGAIKLDDKDPQFKDNVILLNKHIDA
AKZ19095.1       HWPQIAQFAPSASAFFGMSRIGMEVTPSGTWLTYHGAIKLDDKDPQFKDNVILLNKHIDA
AKZ19084.1       HWPQIAQFAPSASAFFGMSRIGMEVTPSGTWLTYHGAIKLDDKDPQFKDNVILLNKHIDA
AIA62318.1       HWPQIAQFAPSASAFFGMSRIGMEVTPSGTWLTYHGAIKLDDKDPQFKDNVILLNKHIDA
AGC74169.1       HWPQIAQFAPSASAFFGMSRIGMEVTPSGTWLTYHGAIKLDDKDPQFKDNVILLNKHIDA
AAZ67039.1       YWPQIAQFAPSASAFFGMSRIGMEVTPLGTWLTYHGAIKLDDKDPQFKDNVILLNKHIDA
Q0Q468.1         YWPQIAQFAPSASAFFGMSRIGMEVTPLGTWLTYHGAIKLDDKDPQFKDNVILLNKHIDA
ARI44802.1       HWPQIAQFAPSASAFFGMSRIGMEVTPSGTWLTYHGAIKLDDKDPQFKDNVILLNKHIDA
ARI44807.1       HWPQIAQFAPSASAFFGMSRIGMEVTPSGTWLTYHGAIKLDDKDPQFKDNVILLNKHIDA
ADE34820.1       HWPQIAQFAPSASAFFGMSRIGMEVTPSGTWLTYHGAIKLDDKDPQFKDNVILLNKHIDA
Q3LZX4.1         HWPQIAQFAPSASAFFGMSRIGMEVTPSGTWLTYHGAIKLDDKDPQFKDNVILLNKHIDA
AAZ41337.1       HWPQIAQFAPSASAFFGMSRIGMEVTPSGTWLTYHGAIKLDDKDPQFKDNVILLNKHIDA
ADE34730.1       HWPQIAQFAPSASAFFGMSRIGMEVTPSGTWLTYHGAIKLDDKDPQFKDNVILLNKHIDA
ADE34787.1       HWPQIAQFAPSASAFFGMSRIGMEVTPSGTWLTYHGAIKLDDKDPQFKDNVILLNKHIDA
ATO98129.1       HWPQIAQFAPSASAFFGMSRIGMEVTPSGTWLTYHGAIKLDDKDPQFKDNVILLNKHIDA
ALK02467.1       HWPQIAQFAPSASAFFGMSRIGMEVTPSGTWLTYHGAIKLDDKDPQFKDNVILLNKHIDA
AGZ48815.1       HWPQIAQFAPSASAFFGMSRIGMEVTPSGTWLTYHGAIKLDDKDPQFKDNVILLNKHIDA
ATO98228.1       HWPQIAQFAPSASAFFGMSRIGMEVTPSGTWLTYHGAIKLDDKDPQFKDSVILLNKHIDA
ATO98190.1       HWPQIAQFAPSASAFFGMSRIGMEVTPSGTWLTYHGAIKLDDKDPQFKDNVILLNKHIDA
ATO98154.1       HWPQIAQFAPSASAFFGMSRIGMEVTPSGTWLTYHGAIKLDDKDPQFKDNVILLNKHIDA
AAR12990.1       HWPQIAQFAPSASAFFGMSRIGMEVTPSGTWLTYHGAIKLDDKDPQFKDNVILLNKHIDA
AAU04673.1       QWPQIAQFAPSASAFFGMSRIGMEVTPSGTWLTYHGAIKLDDKDPQFKDNVILLNKHIDA
ATO98142.1       HWPQIAQFAPSASAFFGMSRIGMEVTLSGTWLTYHGAIKLDDKDPQFKDNVILLNKHIDA
AAP82974.1       HWPQIAQFAPSASAFFGMSRIGMEAAPSGTWLTYHGAIKLDDKDPQFKDNVILLNKHIDA
ATO98117.1       HWPQIAQFAPSASAFFGMSRIGMEVTPSGTWLTYHGAIKLDDKDPQFKDNVILLNKHIDA
AAX16200.1       HWPQIAQFAPSASAFFGMSRIGMEVTPSGTWLTYHGAIKLDDKDPQFKDNVILLNKHIDA
AAV91640.1       QWPQIAQFAPSASAFFGMSRIGMEVTPSGTWLTYHGAIKLDDKDPQFKDNVILLNKHIDA
AAU04642.1       QWPQIAQFAPSASAFFGMSRIGMEVTPSGTWLTYHGAIKLDDKDPQFKDNVILLNKHIDA
AAV49739.1       QWPQIAQFAPSASAFFGMSRIGMEVTPSGTWLTYHGAIKLDDKDPQFKDNVILLNKHIDA
AAS48456.1       HWPQIAQFAPSASAFFGMSRIGMEVTPSGTWLTYHGAIKLDDKDPQFKDNVILLNKHIDA
AAU04658.1       QWPQIAQFAPSASAFFGMSRIGMEVTPSGTWLTYHGAIKLDDKDPQFKDNVILLNKHIDA
ACB69868.1       HWPQIAQFAPSASAFFGMSRIGMEVTPSGTWLTYHGAIKLDDKDPQFKDNVILLNKHIDA
QDF43833.1       YWPQIAQFAPSASAFFGMSRIGMEVTPSGTWLTYHGAIKLDDKDPQFKDNVILLNKHIDA
ACZ72205.1       HWPQIAQFAPSASAFFGMSRIGMEVTPSGTWLTYHGAIKLDDKDPQFKDNVILLNKHIDA
AAP33707.1       HWPQIAQFAPSASAFFGMSRIGMEVTPSGTWLTYHGAIKLDDKDPQFKDNVILLNKHIDA
ACZ72117.1       HWPQIAQFAPSASAFFGMSRIGMEVTPSGTWLTYHGAIKLDDKDPQFKDNVILLNKHIDA
AAT76155.1       HWPQIAQFAPSASAFFGMSRIGMEVTPSGTWLTYHGAIKLDDKDPQFKDNVILLNKHIDA
AGZ48841.1       HWPQIAQFAPSASAFFGMSRIGMEVTPSGTWLTYHGAIKLDDKDPQFKDNVILLNKHIDA
ATO98202.1       HWPQIAQFAPSASAFFGMSRIGMEVTPSGTWLTYHGAIKLDDKDPQFKDNVILLNKHIDA
ATO98166.1       HWPQIAQFAPSASAFFGMSRIGMEVTPSGTWLTYHGAIKLDDKDPQFKDNVILLNKHIDA
ACZ71986.1       HWPQIAQFAPSASAFFGMSRIGMEVTPSGTWLTYHGAIKLDDKDPQFKDNVILLNKHIDA
AAP30714.1       HWPQIAQFAPSASAFFGMSRIGMEVTPSGTWLTYHGAIKLDDKDPQFKDNVILLNKHIDA
AAR87518.1       HWPQIAQFAPSASAFFGMSRIGMEVTPSGTWLTYHGAIKLDDKDPQFKDNVILLNKHIDA
QDF43828.1       HWPQIAQFAPSASAFFGMSRIGMEVTPSGTWLTYHGAIKLDDKDPQFKDNVILLNKHIDA
ATO98178.1       HWPQIAQFAPSASAFFGMSRIGMEVTPSGTWLTYHGAIKLDDKDPQFKDNVILLNKHIDA
NP_828858.1      HWPQIAQFAPSASAFFGMSRIGMEVTPSGTWLTYHGAIKLDDKDPQFKDNVILLNKHIDA
ACU31047.1       HWPQIAQFAPSASAFFGMSRIGMEVTSSGTWLTYHGAIKLDDKDPQFKDNIILLNKHIDA
AID16719.1       HWPQIAQFAPSASAFFGMSRIGMEVTPSGTWLTYHGAIKLDDKDPQFKDNVILLNKHIDA
QDF43818.1       HWPQIAQFAPSASAFFGMSRIGMEVTPSGTWLTYHGAIKLDDKDPQFKDNVMLLNKHIDA
AAZ67043.1       HWPQIAQFAPSASAFFGMSRIGMEVTPSGTWLTYHGAIKLDDKDPQFKDNVILLNKHIDA
Q3I5I7.1         HWPQIAQFAPSASAFFGMSRIGMEVTPSGTWLTYHGAIKLDDKDPQFKDNVILLNKHIDA
ADE34763.1       HWPQIAQFAPSASAFFGMSRIGMEVTPSGTWLTYHGAIKLDDKDPQFKDNVILLNKHIDA
AHX37566.1       HWPQIAQFAPSASAFFGMSRIGMEVTPSGTWLTYHGAIKLDDKDPQFKDNVIMLNKHIDA
ARO76389.1       HWPQIAQFAPSASAFFGMSRIGMEVTPSGTWLTYHGAIKLDDKDPQFKDNVIMLNKHIDA
AGC74175.1       HWPQIAQFAPSASAFFGMSRIGMEVTPSGTWLTYHGAIKLDDKDPQFKDNVILLNKHIDA
AIA62338.1       HWPQIAQFAPSASAFFGMSRIGMEVTPSGTWLTYHGAIKLDDKDPQFKDNVILLNKHIDA
AIA62328.1       HWPQIAQFAPSASAFFGMSRIGMEVTPSGTWLTYHGAIKLDDKDPQFKDNVILLNKHIDA
ACU31039.1       HWPQIAQFAPSASAFFGMSRIGMEVTPSGTWLTYHGAIKLDDKDPQFKDNVILLNKHIDA
QDF43823.1       HWPQIAQFAPSASAFFGMSRIGMEVTPSGTWLTYHGAIKLDDKDPQFKDNVILLNKHIDA
QIA48648.1       HWPQIAQFAPSASAFFGMSRIGMEVTPSGTWLTYTGAIKLDDKDPSFKDNVILLNKHIDA
QIQ54056.1       HWPQIAQFAPSASAFFGMSRIGMEVTPSGTWLTYTGAIKLDDKDPSFKDNVILLNKHIDA
QIA48621.1       HWPQIAQFAPSASAFFGMSRIGMEVTPSGTWLTYTGAIKLDDKDPSFKDNVILLNKHIDA
QIA48630.1       HWPQIAQFAPSASAFFGMSRIGMEVTPSGTWLTYTGAIKLDDKDPSFKDNVILLNKHIDA
QIA48639.1       HWPQIAQFAPSASAFFGMSRIGMEVTPSGTWLTYTGAIKLDDKDPSFKDNVILLNKHIDA
AVP78038.1       HWPQIAQFAPSASAFFGMSRIGMEVTPSGTWLTYHGAIKLDDKDPQFKDNVILLNKHIDA
AVP78049.1       HWPQIAQFAPSASAFFGMSRIGMEVTPSGTWLTYHGAIKLDDKDPQFKDNVILLNKHIDA
QIS61476.1       XXXXXAQFAPSASAFFGMSRIGMEVTPSGTWLTYTGAIKLDDKDPNFKDQVILLNKHIDA
QIQ50109.1       HWPQIAQFAPSASAFFGMSRIGMEVTPSGTWLTYTGAIKLDDKDPNFKDQVILLNKHIDA
QHR63308.1       HWPQIAQFAPSASAFFGMSRIGMEVTPSGTWLTYTGAIKLDDKDPNFKDQVILLNKHIDA
QIS30122.1       HWPQIAQFAPSASAFFGMSRIGMEVTPSGTWLTYTGAIKLDDKDPNFKDQVILLNKHIDA
QIJ96530.1       HWPQIAQFAPSASAFFGMSRIGMEVTPSGTWLTYTGAIKLDDKDPNFKDQVILLNKHIDA
QIQ08827.1       HWPQIAQFAPSASAFFGMSRIGMEVTPSGTWLTYTGAIKLDDKDPNFKDQVILLNKHIDA
QIS30182.1       HWPQIAQFAPSASAFFGMSRIGMEVTPSGTWLTYTGAIKLDDKDPNFKDQVILLNKHIDA
QIH45050.1       HWPQIAQFAPSASAFFGMSRIGMEVTPSGTWLTYTGAIKLDDKVPNFKDQVILLNKHIDA
QIS30511.1       HWPQIAQFAPSASAFFGMSRIGMEVTPSGTWLTYTGAIKLDDKDPNFKDQVILLNKHIDA
QHW06046.1       HWPQIAQFAPSASAFFGMSRIGMEVTPSGTWLTYTGAIKLDDKDPNFKDQVILLNKHIDA
QIM47464.1       HWPQIAQFAPSASAFFGMSRIGMEVTPSGTWLTYTGAIKLDDKDPNFKDQVILLNKHIDA
QIS30192.1       HWPQIAQFAPSASAFFGMSRIGMEVTPSGTWLTYTGAIKLDDKDPNFKDQVILLNKHIDA
QII57305.1       HWPQIAQFAPSASAFFGMSRIGMEVTPSGTWLTYTGAIKLDDKDPNFKDQVILLNKHIDA
QIE07458.1       HWPQIAQFAPSASAFFGMSRIGMEVTPSGTWLTYTGAIKLDDKDSNFKDQVILLNKHIDA
QIS61010.1       HWPQIAQFAPSASAFFGMSRIGMEVTPSGTWLTYTGAIKLDDKDPNFKDQVILLNKHIDA
QIS30062.1       HWPQIAQFAPSASAFFGMSRIGMEVTPSGTWLTYTGAIKLDDKDPNFKDQVILLNKHIDA
QIQ50129.1       HWPQIAQFAPSASAFFGMSRIGMEVTPSGTWLTYTGAIKLDDKDPNFKDQVILLNKHIDA
QIS30552.1       HWPQIAQFAPSASAFFGMSRIGMEVTPSGTWLTYTGAIKLDDKDPNFKDQVILLNKHIDA
QIS60866.1       HWPQIAQFAPSASAFFGMSRIGMEVTPSGTWLTYTGAIKLDDKDPNFKDQVILLNKHIDA
QHO62884.1       HWPQIAQFAPSASAFFGMSRIGMEVTPSGTWLTYTGAIKLDDKDPNFKDQVILLNKHIDA
QIS30502.1       HWPQIAQFAPSASAFFGMSRIGMEVTPSGTWLTYTGAIKLDDKDPNFKDQVILLNKHIDA
QIC50514.1       HWPQIAQFAPSASAFFGMSRIGMEVTPSGTWLTYTGAIKLDDKDPNFKDQVILLNKHIDA
BCB97908.1       HWPQIAQFAPSASAFFGMSRIGMEVTPSGTWLTYTGAIKLDDKDPNFKDQVILLNKHIDA
QIS60878.1       HWPQIAQFAPSASAFFGMSRIGMEVTPSGTWLTYTGAIKLDDKDPNFKDQVILLNKHIDA
YP_009724397.2   HWPQIAQFAPSASAFFGMSRIGMEVTPSGTWLTYTGAIKLDDKDPNFKDQVILLNKHIDA
QIQ49839.1       HWPQIAQFAPSASAFFGMSRIGMEVTPSGTWLTYTGAIKLDDKDPNFKDQVILLNKHIDA
YP_003858591.1   NWPQIAQFAPSASAFFGMSRIGMEVTPTGTWLTYNGAIKLDDKDPNFKDQVILLNKHIDA
APO40586.1       NWPQIAQFAPSASAFFGMSRIGMEVTPTGTWLTYHGAIKLDDKDPNFKDQVILLNKHIDA
```

```
AAS01074.1       ............................................................
ABD75315.1       YKTFPPTEPKKDKKKKTDEAQPLPQRQKKQPTVTLLPAADMDDFSRQLQNSMSGASADST
AIA62298.1       YKTFPPTEPKKDKKKKTDEAQPLPQRQKKQPTVTLLPAADMDDFSRQLQNSMSGASADST
ABG47067.1       YKTFPPTEPKKDKKKKTDEAQPLPQRQKKQPTVTLLPAADMDDFSRQLQNSMSGASADST
ANA96035.1       YKTFPPTEPKKDKKKKTDEAQPLPQRK.KQPTVTLLPAADMDDFSRQLQNSMSGASADST
AIA62285.1       YKTFPPTEPKKDKKKKTDEAQPLPQRK.KQPTVTLLPAADMDDFSRQLQNSMSGASADST
ASO66816.1       YKTFPPTEPKKDKKKKTDEAQPLPQRK.KLPTVTLLPAADMDDFSRQLQNSMSGASADST
AKZ19095.1       YKTFPPTEPKKDKKKKTDEAQPLPQRQKKQPTVTLLPAADMDDFSRQLQNSMSEASADST
AKZ19084.1       YKTFPPTEPKKDKKKKTDEAQPLPQRQKKQPTVTLLPAADMDDFSRQLQNSMSGASADST
AIA62318.1       YKTFPPTEPKKDKKKKTDEAQPLPQRK.KQPTVTLLPAADMDDFSRQLQNSMSGASADST
AGC74169.1       YKTFPPTEPKKDKKKKTDEAQPLPQRK.KQPTVTLLPAADMDDFSRQLQNSMSGASADST
AAZ67039.1       YKTFPPTEPKKDKKKKTDEAQPLPQRK.KQPTVTLLPAADMDDFSRQLQNSMSGASADST
Q0Q468.1         YKAFPPTEPKKDKKKKTDEAQPLPQRK.KQPTVTLLPAADMDDFSRQLQNSMSGASADST
ARI44802.1       YKTFPPTEPKKDKKKKTDEAQPLPQRQKKQPIVTLLPAADMDDFSRQLQNSMSGASADST
ARI44807.1       YKTFPPTEPKKDKKKKTDEAQPLPQRQKKQPIVTLLPAADMDDFSRQLQNSMSGASADST
ADE34820.1       YKTFPPTEPKKDKKKKTDEAQPLPQRQKKQPTVTLLPAADMDDFSRQLQHSMSGASADST
Q3LZX4.1         YKTFPPTEPKKDKKKKTDEAQPLPQRQKKQPTVTLLPAADMDDFSRQLQHSMSGASADST
AAZ41337.1       YKTFPPTEPKKDKKKKTDEAQPLPQRQKKQPTVTLLPAADMDDFSRQLQHSMSGASADST
ADE34730.1       YKTFPPTEPKKDKKKKTDEAQPLPQRQKKQPTVTLLPAADMDDFSRQLQHSMSGASADST
ADE34787.1       YKTFPPTEPKKDKKKKTDEAQPLPQRQKKQPTVTLLPAADMDDFSRQLQHSMSGASADST
ATO98129.1       YKTFPPTEPKKDKKKKTDEAQPLPQRQKKQPTVTLLPAADMDDFSRQLQNSMSGASADST
ALK02467.1       YKTFPPTEPKKDKKKKTDEAQPLPQRQKKQPTVTLLPAADMDDFSRQLQNSMSGASADST
AGZ48815.1       YKTFPPTEPKKDKKKKTDEAQPLPQRQKKQPTVTLLPAADMDDFSRQLQNSMSGASADST
ATO98228.1       YKTFPPTEPKKDKKKKTDEAQPLPQRQKKQPTVTLLPAADMDDFSRQLQNSMSGASADST
ATO98190.1       YKTFPPTEPKKDKKKKTDEAQPLPQRQKKQPTVTLLPAADMDDFSRQLQNSMSGASADST
ATO98154.1       YKTFPPTEPKKDKKKKTDEAQPLPQRQKKQPTVTLLPAADMDDFSRQLQSSMSGASADST
AAR12990.1       YKTFPPTEPKKDKKKKTDEAQPLPQRQKKQPTVTLLPAADMDDFSRQLQNSMSGASADST
AAU04673.1       YKTFPPTEPKKDKKKKTDEAQPLPQRQKKQPTVTLLPAADMDDFSRQLQNSMSGASADST
ATO98142.1       YKTFPPTEPKKDKKKKTDEAQPLPQRQKKQPTVTLLPAADMDDFSRQLQNSMSGASADST
AAP82974.1       YKTFPPTEPKKDKKKKTDEAQPLPQRQKKQPTVTLLPAADMDDFSRQLQNSMSGASADST
ATO98117.1       YKTFPPTEPKKDKKKKTDEAQPLPQRQKKQPIVTLLPAADMDDFSRQLQNSMSGASADST
AAX16200.1       YKTFPPTEPKKDKKKKTDEAQPLPQRQKKQPTVTLLPAADMDDFSRQLQNSMSGASADST
AAV91640.1       YKTFPPTEPKKDKKKKTDEAQPLLQRQKKQPTVTLLPAADMDDFSRQLQNSMSGASADST
AAU04642.1       YKTFPPTEPKKDKKKKTDEAQPLLQRQKKQPTVTLLPAADMDDFSRQLQNSMSGASADST
AAV49739.1       YKTFPPTEPKKDKKKKTDEAQPLPQRQKKQPTVTLLPAADMDDFSRQLQNSMSGASADST
AAS48456.1       YKTFPPTEPKKDKKKKTDEAQPLPQRQKKQPTVTPLPAADMDDFSRQLQNSMSGASADST
AAU04658.1       YKTFPPTEPKKDKKKKTDEAQPLPQRQKKQPTVTLLPAADMDDFSRQLQNSMSGASADST
ACB69868.1       YKTFPPTEPKKDKKKKTDEAQPLPQRQKKQPTVTLLPAADMDDFSRQLQNSMSGASADST
QDF43833.1       YKTFPPTEPKKDKKKKTDEAQPLPQRQKKQPTVTLLPAADMDDFSRQLQNSMSGASADST
ACZ72205.1       YKTFPPTEPKKDKKKKTDEAQPLPQRQKKQPTVTLLPAADMDDFSRQLQNSMSGASADST
AAP33707.1       YKTFPPTEPKKDKKKKTDEAQPLPQRQKKQPTVTLLPAADMDDFSRQLQNSMSGASADST
ACZ72117.1       YKTFPPTEPKKDKKKKTDEAQPLPQRQKKQPTVTLLPAADMGDFSRQLQNSMSGASADST
AAT76155.1       YKTFPPTEPKKDKKKKTDEAQPLPQRQKKQPTVTLLPAADMDDFSRQLQNSMSGASADST
AGZ48841.1       YKTFPPTEPKKDKKKKTDEAQPLPQRQKKQPTVTLLPAADMDDFSRQLQNSMSGASADST
ATO98202.1       YKTFPPTEPKKDKKKKTDEAQPLPQRQKKQPTVTLLPAADMDDFSRQLQNSMSGASADST
ATO98166.1       YKTFPPTEPKKDKKKKTDEAQPLPQRQKKQPTVTLLPAADMDDFSRQLQNSMSGASADST
ACZ71986.1       YKTFPPTEPKKDKKKKTDEAQPLPQRQKKQPTVTLLPAADMDDFSRQLQNSMSGASADST
AAP30714.1       YKTFPPTEPKKDKKKKTDEAQPLPQRQKKQPTVTLLPAADMDDFSRQLQNSMSGASADST
AAR87518.1       YKTFPPTEPKKDKKKKTDEAQPLPQRQKKQPTVTLLPAADMDDFSRQLQNSMSGASADST
QDF43828.1       YKTFPPTEPKKDKKKKTDEAQPLPQRQKKQPTVTLLPAADMDDFSRQLQNSMSGASADST
ATO98178.1       YKTFPPTEPKKDKKKKTDEAQPLPQRQKKQPTVTLLPAADMDDFSRQLQNSMSGASADST
NP_828858.1      YKTFPPTEPKKDKKKKTDEAQPLPQRQKKQPTVTLLPAADMDDFSRQLQNSMSGASADST
ACU31047.1       YKTFPPTEPKKDKKKKTDEAQPLPQRPKKQPTVTLLPAADMDDFSRQLQNSMSGASADST
AID16719.1       YKTFPPTEPKKDKKKKTDEAQPLPQRPKKQPTVTLLPAADMDDFSRQLQNSMSGASADST
QDF43818.1       YKAFPPTEPKKDKKKKTDEAQPLPQRQKKQPTVTLLPAADMDDFSRQLQNSMSGASADST
AAZ67043.1       YKTFPPTEPKKDKKKKTDEAQPLPQRQKKQPTVTLLPAADMDDFSRQLQNSMSGASADST
Q3I5I7.1         YKIFPPTEPKKDKKKKTDEAQPLPQRQKKQPTVTLLPAADMDDFSRQLQNSMSGASADST
ADE34763.1       YKTFPPTEPKKDKKKKTDEAQPLPQRQKKQPTVTLLPAADMDDFSRQLQKSMSGASADST
AHX37566.1       YKTFPPTEPKKDKKKKTDEAQPLPQRQKKQPTVTLLPAADMDDFSRQLQNSMSGASADST
ARO76389.1       YKTFPPTEPKKDKKKKTDEAQPLPQRQKKQHTVTLLPAADMDDFSRQLQNSMSGASADST
AGC74175.1       YKTFPPTEPKKDKKKKTDEAQPLPQRQKKQPTVTLLPAADMDDFSRQLQNSMSGASADST
AIA62338.1       YKTFPPTEPKKDKKKKTDEAQPLPQRQKKQPTVTLLPAADMDDFSRQLQNSMGGASADST
AIA62328.1       YKTFPPTEPKKDKKKKTDEAQPLPQRQKKQPTVTLLPAADMDDFSRQLQNSMSGASADST
ACU31039.1       YKTFPPTEPKKDKKKKTDEAQPLPQRQKKQPTVTLLPAADMDDFSRQLQNSMSGASADST
QDF43823.1       YKTFPPTEPKKDKKKKTDEAQPLPQRQKKQPTVTLLPAADMDDFSRQLQNSMSGASADST
QIA48648.1       YKTFPPTEPKKDKKKKTDESQPLPQRQKKQQTVTLLPAADLDDFSKQLQQSMSSADSTQA
QIQ54056.1       YKTFPPTEPKKDKKKKTDESQPLPQRQKKQQTVTLLPAADLDDFSKQLQQSMSSADSTQA
QIA48621.1       YKTFPPTEPKKDKKKKTDESQPLPQRQKKQQTVTRLPAADLDDFSKQLQQSMSSADSTQA
QIA48630.1       YKTFPPTEPKKDKKKKTDESQPLPQRQKKQQTVTLLPAADLDDFSKQLQQSMSSADSTQA
QIA48639.1       YKTFPPTEPKKDKKKKTDESQPLPQRQKKQQTVTLLPAADLDDFSKQLQQSMSSADSTQA
AVP78038.1       YKTFPPTEPKKDKKKKADELQALPQRQKKQQTVTLLPAADLDEFSKQLQQSMSGTDSTQA
AVP78049.1       YKTFPPTEPKKDKKKKADELQALPQRQKKQQTVTLLPAADLDEFSKQLQQSMSGTDSTQA
QIS61476.1       YKTFPPTEPKKDKKKKADETQALPQRQKKQQTVTLLPAADLDDFSKQLQQSMSSADSTQA
QIQ50109.1       YKTFPPTEPKKDKKKKADETQALPQRQKKQQTVTLLPAADLDDFSKQLQQSMSSADSTQA
QHR63308.1       YKTFPPTEPKKDKKKKADETQALPQRQKKQQTVTLLPAADLDDFSKQLQQSMSSADSTQA
QIS30122.1       YKTXXPTEPKKDKKKKADETQALPQRQKKQQTVTLLPAADLDDFSKQLQQSMSSADSTQA
QIJ96530.1       YKTFPPTEPKKDKKKKADETQALPQRQKKQQTVTLLPAADLDDFSKQLQQSMSSADSTQA
QIQ08827.1       YKTFPPTEPKKDKKKKADETQALPQRQKKQQTVTLLPAADLDDFSKQLQQSMSSADSTQA
QIS30182.1       YKTFPPTEPKKDKKKKADETQALPQRQKKQQTVTLLPAADLDDFSKQLQQSMSSADSTQA
QIH45050.1       YKTFPPTEPKKDKKKKADETQALPQRQKKQQTVTLLPAADLDDFSKQLQQSMSSADSTQA
QIS30511.1       YKTFPXTEPKKDKKKKADETQALPQRQKKQQTVTLLPAADLDDFSKQLQQSMSSADSTQA
QHW06046.1       YKTFPPTEPKKDKKKKADETQALPQRQKKQQTVTLLPAADLDDFSKQLQQSMSSADSTQA
QIM47464.1       YKTFPPTEPKKDKKKKADETQALPQRQKKQQTVTLLPAADLDDFSKQLQQSMSSADSTQA
QIS30192.1       YKTFPPTXPKKDKKKKADETQALPQRQKKQQTVTLLPAADLDDFSKQLQQSMSSADSTQA
QII57305.1       YKTFPPTEPKKDKKKKADETQALPQRQKKQQTVTLLPAADLDDFSKQLQQSMSSADSTQA
QIE07458.1       YKTFPPTEPKKDKKKKADETQALPQRQKKQQTVTLLPAADLDDFSKQLQQSMSSADSTQA
QIS61010.1       YKTFPPTEPKKDKKKKADETQALPQRQKKQQTVTLLPAADLDDFSKQLQQSMSSADSTQA
QIS30062.1       YKTFPPTEPKKDKKKKADETQALPQRQKKQQTVTLLPAADLDDFSKQLQQSMSSADSTQA
QIQ50129.1       YKTFPPTEPKKDKKKKADETQALPQRQKKQQTVTLLPAADLDDFSKQLQQSMSSADSTQA
QIS30552.1       YKTFPPTEPKKDKKKKADETQALPQRQKKQQTVTLLPAADLDDFSKQLQQSMSSADSTQA
QIS60866.1       YKTFPPTEPKKDKKKKADETQALPQRQKKQQTVTLLPAADLDDFSKQLQQSMSSADSTQA
QHO62884.1       YKTFPPTEPKKDKKKKADETQALPQRQKKQQTVTLLPAADLDDFSKQLQQSMSSADSTQA
QIS30502.1       YKTFPPTEPKKDKKKKADETQALPQRQKKQQTVTLLPAADLDDFSKQLQQSMSSADSTQA
QIC50514.1       YKTFPPTEPKKDKKKKADETQALPQRQKKQQTVTLLPAADLDDFSKQLQQSMSSADSTQA
BCB97908.1       YKTFPPTEPKKDKKKKADETQALPQRQKKQQTVTLLPAADLDDFSKQLQQSMSSADSTQA
QIS60878.1       YKTFPPTEPKKDKKKKADETQALPQRQKKQQTVTLLPAADLDDFSKQLQQSMSSADSTQA
YP_009724397.2   YKTFPPTEPKKDKKKKADETQALPQRQKKQQTVTLLPAADLDDFSKQLQQSMSSADSTQA
QIQ49839.1       YKTFPPTEPKKDKKKKADETQALPQRQKKQQTVTLLPAADLDDFSKQLQQSMSSADSTQA
YP_003858591.1   YKTFPPTEPKKDKKKKADEVQSLPQRQKKQATVTLLPAADLDDFSKQLQNSMNASPDSTQ
APO40586.1       YKTFPPTEPKKDKKKKADEVQPLPQRQKKQPTVTLLPAAELDDFSKQLQDSMNGASDSTQ
```

| | |
|---|---|
| AAS01074.1 | . . |
| ABD75315.1 | Q A |
| AIA62298.1 | Q A |
| ABG47067.1 | Q A |
| ANA96035.1 | Q A |
| AIA62285.1 | Q A |
| ASO66816.1 | Q A |
| AKZ19095.1 | Q A |
| AKZ19084.1 | Q A |
| AIA62318.1 | Q A |
| AGC74169.1 | Q A |
| AAZ67039.1 | Q A |
| Q0Q468.1 | Q A |
| ARI44802.1 | Q A |
| ARI44807.1 | Q A |
| ADE34820.1 | Q A |
| Q3LZX4.1 | Q A |
| AAZ41337.1 | Q A |
| ADE34730.1 | Q A |
| ADE34787.1 | Q A |
| ATO98129.1 | Q A |
| ALK02467.1 | Q A |
| AGZ48815.1 | Q A |
| ATO98228.1 | Q A |
| ATO98190.1 | Q A |
| ATO98154.1 | Q A |
| AAR12990.1 | Q A |
| AAU04673.1 | Q A |
| ATO98142.1 | Q A |
| AAP82974.1 | Q A |
| ATO98117.1 | Q A |
| AAX16200.1 | Q A |
| AAV91640.1 | Q A |
| AAU04642.1 | Q A |
| AAV49739.1 | Q A |
| AAS48456.1 | Q A |
| AAU04658.1 | Q A |
| ACB69868.1 | Q A |
| QDF43833.1 | Q A |
| ACZ72205.1 | Q A |
| AAP33707.1 | Q A |
| ACZ72117.1 | Q A |
| AAT76155.1 | Q A |
| AGZ48841.1 | Q A |
| ATO98202.1 | Q A |
| ATO98166.1 | Q A |
| ACZ71986.1 | Q A |
| AAP30714.1 | Q A |
| AAR87518.1 | Q A |
| QDF43828.1 | Q A |
| ATO98178.1 | Q A |
| NP_828858.1 | Q A |
| ACU31047.1 | Q A |
| AID16719.1 | Q A |
| QDF43818.1 | Q A |
| AAZ67043.1 | Q A |
| Q3I5I7.1 | Q A |
| ADE34763.1 | Q A |
| AHX37566.1 | Q A |
| ARO76389.1 | Q A |
| AGC74175.1 | Q A |
| AIA62338.1 | Q A |
| AIA62328.1 | Q A |
| ACU31039.1 | Q A |
| QDF43823.1 | Q A |
| QIA48648.1 | . . |
| QIQ54056.1 | . . |
| QIA48621.1 | . . |
| QIA48630.1 | . . |
| QIA48639.1 | . . |
| AVP78038.1 | . . |
| AVP78049.1 | . . |
| QIS61476.1 | . . |
| QIQ50109.1 | . . |
| QHR63308.1 | . . |
| QIS30122.1 | . . |
| QIJ96530.1 | . . |
| QIQ08827.1 | . . |
| QIS30182.1 | . . |
| QIH45050.1 | . . |
| QIS30511.1 | . . |
| QHW06046.1 | . . |
| QIM47464.1 | . . |
| QIS30192.1 | . . |
| QII57305.1 | . . |
| QIE07458.1 | . . |
| QIS61010.1 | . . |
| QIS30062.1 | . . |
| QIQ50129.1 | . . |
| QIS30552.1 | . . |
| QIS60866.1 | . . |
| QHO62884.1 | . . |
| QIS30502.1 | . . |
| QIC50514.1 | . . |
| BCB97908.1 | . . |
| QIS60878.1 | . . |
| YP_009724397.2 | . . |
| QIQ49839.1 | . . |
| YP_003858591.1 | A . |
| APO40586.1 | A . |
